# Cex1 is a new component of the COPI Golgi-to-vacuole intracellular trafficking machinery

**DOI:** 10.1101/414367

**Authors:** Ludovic Enkler, Johann Owen de Craene, Bruno Rinaldi, Philippe Hammann, Osamu Nureki, Bruno Senger, Sylvie Friant, Hubert D. Becker

## Abstract

During translational elongation, aminoacylated tRNA are supplied to the ribosome by RNA-interacting proteins. Cex1, a HEAT-containing protein, has been shown to participate to this tRNA channeling by interacting with aminoacylated tRNA during their export from the nucleus. Here, we show that Cex1 is a component of COPI (coatomer complex I) coated vesicles involved in the Golgi-to-vacuole trafficking pathway in *Saccharomyces cerevisiae*. Cex1 interacts with key components of the COPI coat Sec27, Sec28 and Sec33 proteins. Moreover, fluorescent microscopy indicates that Cex1 is not localized at the nuclear periphery as expected for an effector of nuclear tRNA channeling, but is observed on endosomal trans-Golgi network (TGN) positive structures. This localization relies on the vacuolar protein sorting receptor Vps10. Our data not only resolve the functional ambiguity regarding Cex1 homologues across species, but also point to the possibility to develop yeast-based models to study neurodegenerative disorders linked to the Human Cex1 homologue SCYL1.

## Introduction

In eukaryotes, transcription and translation are physically separated by the nuclear envelope. Transport of macromolecules such as proteins or RNAs between the nucleus and the cytosol is achieved through nuclear pores that assemble into large complexes known as nuclear pore complexes (or NPCs). Molecules smaller than 40 kDa diffuse through the NPCs, while larger molecules need specific import/export machineries. Nuclear import is generally mediated by importins and export by exportins, both belonging to the β-karyopherin family. In the yeast *Saccharomyces cerevisiae*, 14 differents β-karyopherins were identified based on the homology of their i) N-terminus with the Gsp1 GTPase binding domain (RanGTP in mammals) and ii) their C-terminus containing the HEAT (Huntingtin, elongation factor 3 (EF3), protein phosphatase 2A (PP2A), and the yeast kinase TOR1) domain made of repeated motifs (Yoshimura & Hirano, 2016).

In the nucleus of yeast cells, transfer RNAs (tRNAs) are subjected to several modifications such as 5’ and 3’ trimming, base modifications and the addition of the 3′-CCA terminus (Hopper, 2013). Intron-containing tRNAs are exported *via* Los1 to the cytoplasm where they are spliced, modified and finally used for translation (Huh *et al*, 2003; Bertrand *et al*, 1998). Indeed, aminoacylated tRNAs (aa-tRNAs) are delivered to the nuclear receptors Los1 and Msn5 by the nucleolar protein Utp8, (Izaurralde *et al*, 1997; Richards *et al*, 1997; Steiner-Mosonyi *et al*, 2003). Upon export, aa-tRNAs are associated to the elongation factor eEF-1A and participate in cytosolic translation. Different studies have shown that aa-tRNAs can be re-imported into the nucleus by the β-karyopherin Mtr10 in a retrograde transport (Yoshihisa *et al*, 2003; Stanford *et al*, 2004). This nuclear accumulation of aa-tRNAs was observed during amino acids starvation (Shaheen & Hopper, 2005), it has also been proposed that this retrograde transport might be constitutive (Chafe *et al*, 2011). Following this, aa-tRNAs are re-exported from the nucleus *via* several redundant pathways, some still poorly understood.

Cex1 has been identified as a new cytoplasmic component of the tRNA export machinery, based on a yeast triple hybrid assay aimed at identifying protein-RNA interactions (McGuire & Mangroo, 2007). Its implication in tRNA nuclear export was supported by TAP (tandem affinity purification) and pull-down experiments in which Cex1 co-purified with Los1, Gsp1, eEF-1A and the nucleoporin Nup116 (McGuire & Mangroo, 2007). Moreover, the *cex1*Δ *los1*Δ yeast mutant cells were impaired in nuclear tRNA export but cell growth was not affected (McGuire & Mangroo, 2007). In contrast, previous studies showed that cells bearing deletions of *LOS1* and *ARC1* (a cytoplasmic tRNA export protein) genes are not viable (Simos *et al*, 1996). These data suggest that Cex1 function in tRNA nuclear export is distinct from that of Arc1. Moreover, a recent study revealed that Cex1 activates Gsp1 GTPase activity by recruiting the GTPase-activating protein (GAP) Ran1 (McGuire & Mangroo, 2012). This suggests that Cex1 is implicated in the nuclear re-export of aa-tRNAs, with aa-tRNAs being handed from Los1 to eEF1 *via* Cex1 on the cytoplasmic side of the NPC. It was also proposed that Cex1-dependent recruitment of Ran1 at the NPC enables the dissociation of the export complex made of Los1·aa-tRNA and Gsp1. These functions were also proposed to be conserved in mammalian cells (Chafe *et al*, 2011). However, a recent attempt could not reproduce the *cex1*Δ *arc1*Δ lethality casting doubt on the existence of a genetic interaction between *ARC1* and *CEX1* (Johnstone *et al*, 2011), which had also not been found in a *S. cerevisiae* genome-wide synthetic lethality.

We thus re-analyzed the implication of the genetic interaction between *CEX1* and *ARC1* and the participation of Cex1 to nuclear export of aa-tRNAs. By following an unbiased approach, our data unambiguously demonstrate that *CEX1* is not synthetic lethal with *ARC1* suggesting that the participation of Cex1 to nuclear-cytoplasmic transport of aa-tRNAs might as well have been misattributed. Indeed, here we demonstrate that Cex1 is associated to subunits of the coatomer COPI complex by interacting with Sec27, Sec28 and Sec33 COPIb proteins that participates to protein sorting. Our data further identify *CEX1* as a new regulator of the vesicular trafficking from the Golgi to the vacuole, shedding a new light on the cellular function of Cex1 in yeast and on COPI-mediated trafficking. Interestingly, these findings corroborate what is known in mammals, where *CEX1* homologue *SCYL1* is also involved in vesicular complexes and regulates Golgi-to-ER transport both in Human and mice *via* its interaction with COPI (Coatomer complex I) coat proteins (Burman *et al*, 2008; Burman *et al*, 2010).

## Results

### *ARC1* and *CEX1* are not synthetic lethal

To re-analyze the genetic interaction between *ARC1* and *CEX1*, we generated and studied the *cex1*Δ and *arc1*Δ *cex1*Δ deletion strains. Since it was previously shown that the *arc1*Δ *cex1*Δ double deletion was lethal, we first generated a *cex1*Δ strain expressing *CEX1* under its own promoter from a centromeric pRS316 plasmid. Next, the *ARC1* gene was deleted by homologous recombination and Arc1 was not detectable in this strain by an anti-Arc1 western-blot (Fig 1A). To test the genetic interaction, we grew the *arc1*Δ *cex1*Δ cells bearing the pRS316-CEX1 plasmid on 5-FOA plates to induce plasmid loss. After 3 days of incubation at 30°C, several isolated colonies were observed on the 5-FOA plates suggesting that in our genetic background, *ARC1* and *CEX1* are not synthetically lethal. To further confirm these results, a phenotypic analysis of the different strains was done on different selective media. *CEX1* deletion was confirmed by growth of the strain on YPD+G418, *ARC1* deletion was confirmed by growth of the strain on SC-His and finally loss of the pRS316-CEX1 plasmid was confirmed by the absence of growth on SC-Ura (Fig 1B). Since Arc1 was previously associated to the diauxic shift from fermentation to respiration conditions (Frechin et al., 2009; Frechin et al., 2014), we also tested growth on respiration medium (SC-Gly), in presence of sucrose (SC-Suc) as the sole source of carbon, or upon an osmotic shock (1M sorbitol) at different temperatures. The *cex1*Δ *arc1*Δ strain grew like a wild-type (WT) in presence of sucrose (SC-Suc) as the sole source of carbon, or upon an osmotic shock (1M sorbitol; Fig EV1A). The *cex1*Δ *arc1*Δ strain displayed reduced growth on glycerol medium (Fig 1C), a respiratory defect most probably due to *ARC1* deletion since the *arc1*Δ cells showed a similar growth delay (Fig 1C). To ascertain that Cex1 is not only genetically but also functionally independent of Arc1, we tested another *arc1*Δ phenotype, which is vacuolar aa storage and signaling. Arc1 has been shown to bind with a higher affinity several tRNA species and notably tRNA^Pro^, tRNA^Arg^ for which the cognate aa are stored in vacuoles, the compartments responsible for aa storage (Fig EV2B) (Kitamoto *et al*, 1988; Deinert *et al*, 2001). To analyze these vacuolar- and Arc1-dependent functions, growth of the *cex1*Δ, *arc1*Δ and *cex1*Δ *arc1*Δ strains was tested in the presence of molar excess of proline (Pro), arginine (Arg) or isoleucine (Ile) aa (Figure EV2C). None of these conditions affected the survival of the strains (Figure EV2C), indicating that Cex1 does not impact the vacuolar aa storage function. These lines of experiments not only show that *ARC1* and *CEX1* are not genetically linked, but that *CEX1* is not involved in the selection of the carbon (fermentation versus respiration) or aa source during growth of *S. cerevisiae*.

**Figure 1:**
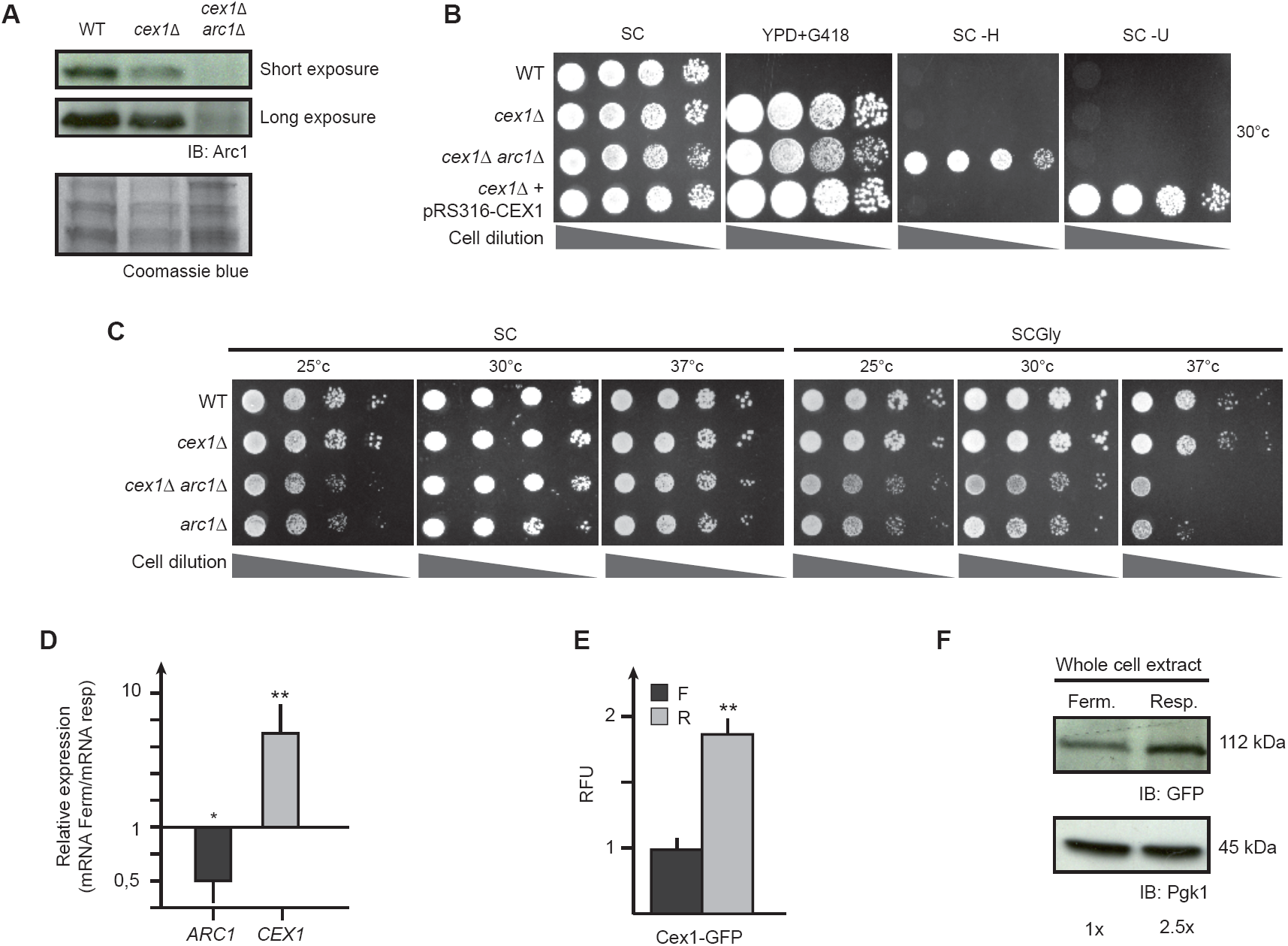
In *Saccharomyces cerevisiae CEX1* and *ARC1* are not genetically linked. **A** Immunodetection of Arc1 in a WT (BY4742), *cex1Δ* and *cex1Δarc1Δ* strains. Coomassie blue staining was also performed to highlight protein levels in each lane. **B** Drop test of the WT, *cex1Δ, cex1Δarc1Δ* and *cex1Δ*+pRS316-*CEX1* strains on rich media in the presence of Geneticin (YPD+G418), and on synthetic selective media (SC-Glc, SC-H, SC-U). Growth was performed at 30 °C for 2 days. **C** Drop test of the WT, *cex1Δ, cex1Δarc1Δ* and *arc1Δ* strains grown for 2 days in fermentation (SC-Glc) or 3 days in respiration (SC-Gly) at 25, 30 and 37 °C. **D** Relative amounts of *ARC1* and *CEX1* mRNAs measured by RT-qPCR in respiration as compared to fermentation. * *p=* 0,005; ** *p*< 0,001. **E** Fluorescence of Cex1-GFP cells grown in fermentation (grey) and respiration (orange) measured by epifluorescence microscopy. At least 5 fields were used for calculation in each condition (n >100 cells). Results are shown in Relative Fluorescence Unit (RFU). ** *p*< 0,001. **F** Immunodetection of Cex1-GFP in whole cell extracts of yeast grown in fermentation or respiration. Pgk1p was used as loading control. Cex1 quantitation is the result of three experiments.

However, when we compared *ARC1* and *CEX1* expressions in fermentation (grown on glucose) and respiration (grown on glycerol) conditions on wild-type yeast cells by reverse transcription quantitative PCR (RT-qPCR), we found that *ARC1* was 0.5-fold downregulated in respiration while *CEX1* was 6-fold upregulated (Fig 1D). This increased expression level was confirmed at the protein level, both by epifluorescence microscopy and immunodetection, where we observed a 1.8- and 2.5-fold increase of Cex1 protein level in respiration compared to fermentation (Fig 1E-F respectively). This would suggest that Cex1 could take over Arc1’s role in exporting tRNA from the nucleus only in respiring yeast cells but not in fermenting ones. However, compared to the wild-type, the *cex1*Δ *arc1*Δ strain did not display any growth defect at 25-30°C on Glycerol medium which excludes an involvement of Cex1 in tRNA nuclear export specifically during respiration. None the less, the expression profile of *CEX1* suggests that Cex1 role is probably more important during respiration than fermentation.

### Cex1 interacts with COPI vesicles coat proteins

To better understand the cellular role of Cex1, we decided to identify its protein partners. Immunoprecipitation (IP) of HA tagged Cex1 proteins coupled to mass-spectrometry analysis was performed on a *cex1*Δ strain bearing the pHAC111-CEX1 plasmid, allowing expression of Cex1-HA under the dependence of its own promoter (Table 1 and 3). Cells were grown in respiratory conditions (SC-Gly) and lysed in the presence of mild detergent to retrieve potential membrane-associated complexes. A closer look at the most detected candidates based on mass-spectrometry spectral counts revealed that most of them belong to the COPI vesicle family, namely: Sec21, Sec26, Sec27, Sec28 and Sec33 (Gabriely *et al*, 2006) (Fig 2A, also see Extended Table 1). Cex1 also interacts with the Golgi protein Erv46, and Sec18 a component of the V- and t-SNARE dissociation complex, although these proteins might bind non-specifically to Cex1-HA given the detection of 3 or 1 spectral counts of these two proteins in the negative MYC-IP controls (Fig 2A;(Shibuya *et al*, 2015)). In this Cex1-HA IP experiment, Epo1 involved in septin-ER tethering, Mlp1 a protein from the nuclear envelope and two cytoplasmic proteins (Bat2 and Tps1) were also detected. We did not anticipate that Cex1, identified as a tRNA-binding protein involved in the nuclear-cytoplasmic export pathway (McGuire & Mangroo, 2007; 2012), would be physically linked to membrane-related proteins involved in intracellular trafficking. To further assess the connection between Cex1 and vesicular trafficking, we looked for evidences of such interactions in the literature. A recent genetic interaction screen done in *S. cerevisiae* showed that 16 genes are genetically linked to *CEX1* and 8 among them encode Golgi and/or COPI vesicle coat proteins (Usaj *et al*, 2017). Moreover, *SEC33* and *SEC26* were also found to be genetically linked to *CEX1* by Usaj et al., 2016 (circled names, Fig 2B). Interestingly, SCYL1 the mammalian homologue of Cex1 also interacts with human COPI components, COPA (Sec33), COPB1 (Sec26) and COPB2 (Sec27) (Ioannou & Summerfeldt, 2015) (Fig 2B). Cellular component GO term analysis of Cex1 interactants revealed that proteins participating in microtubule organizing center part, in cytoplasmic vesicle’s membranes and integral component of organelle’s membrane were enriched in our interaction experiment respectively at 47%, 34% and 22% (Fig 2C). Furthermore, based on the Saccharomyces Genome Database (SGD) and BioGRID3.4, *CEX1* has 15 known physical interactants, and among them only 4 were in our interactome analysis. Interestingly, none of them are related to the intracellular trafficking, pointing toward the importance of the purification conditions that one uses for Cex1 (Fig 2D). Using the same databases, we retrieved 90 genetic interactants, a third (37) being linked to the Golgi and involved in intracellular trafficking. Almost half (15) of these genetic interactants were also retrieved as physical partners in our IP experiments using Cex1-HA as bait (Fig 2E).

**Table 1:**
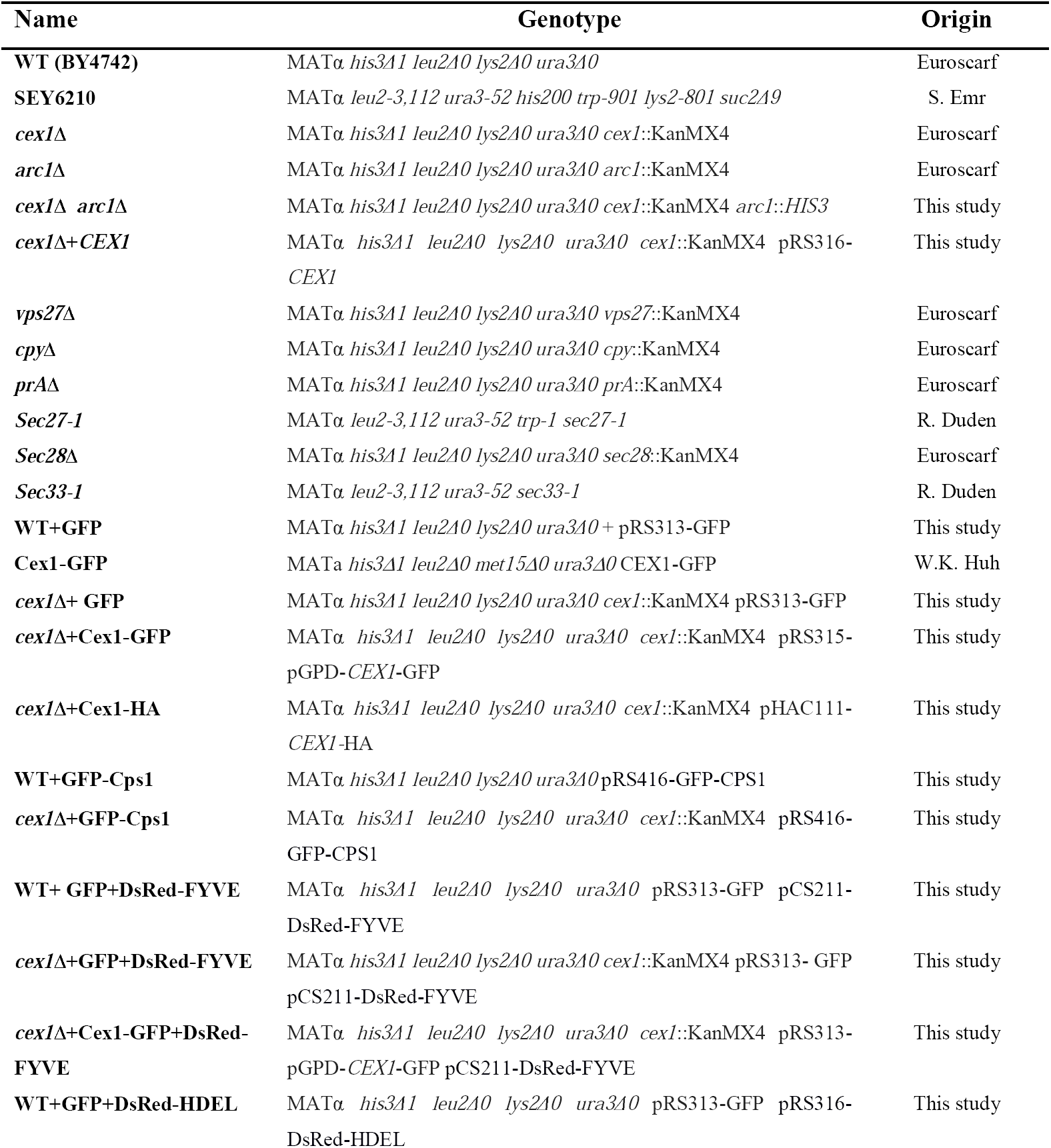
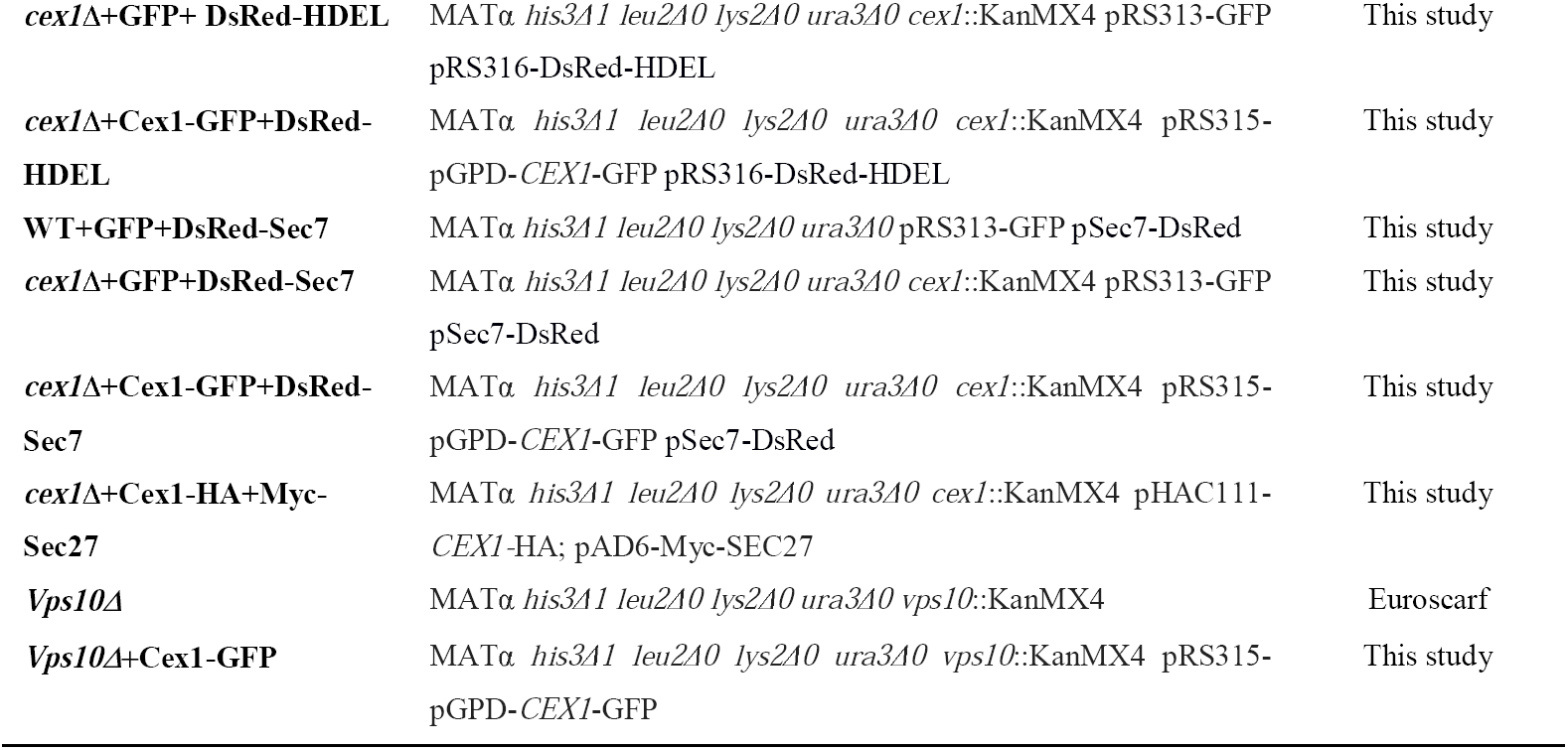
Strains used in this study

**Table 2:**
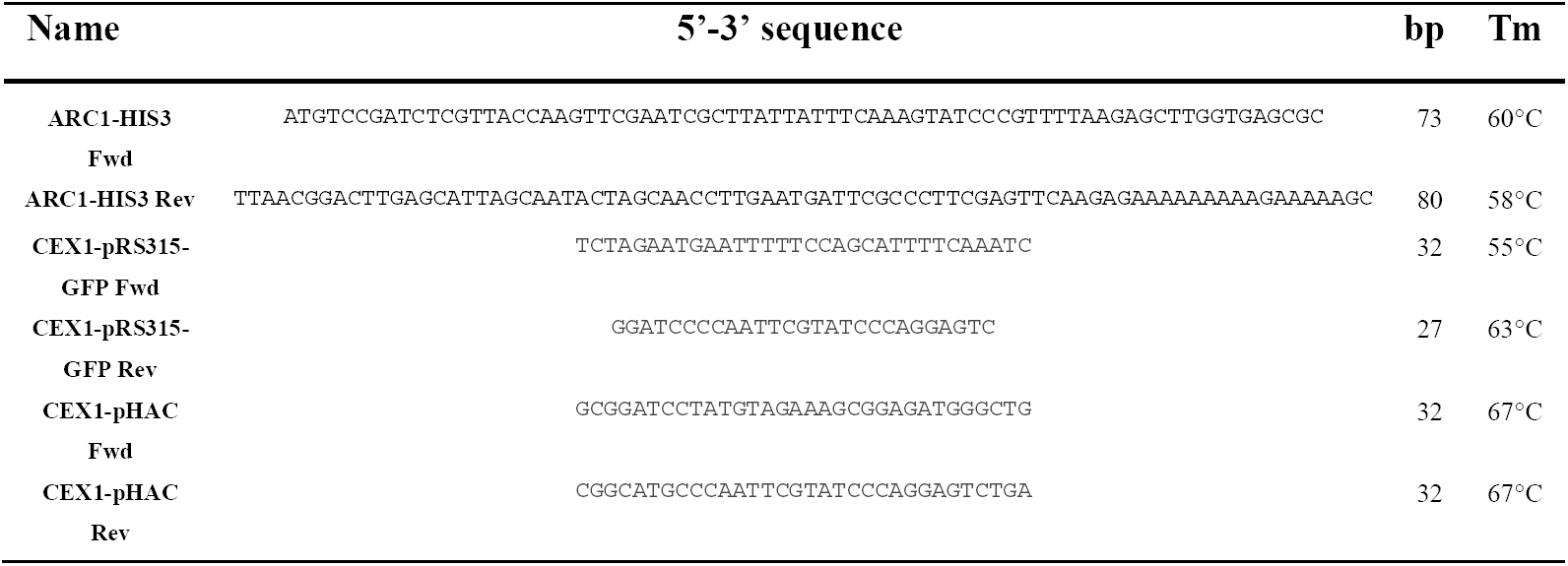
Primers used for cloning

**Table 3:**
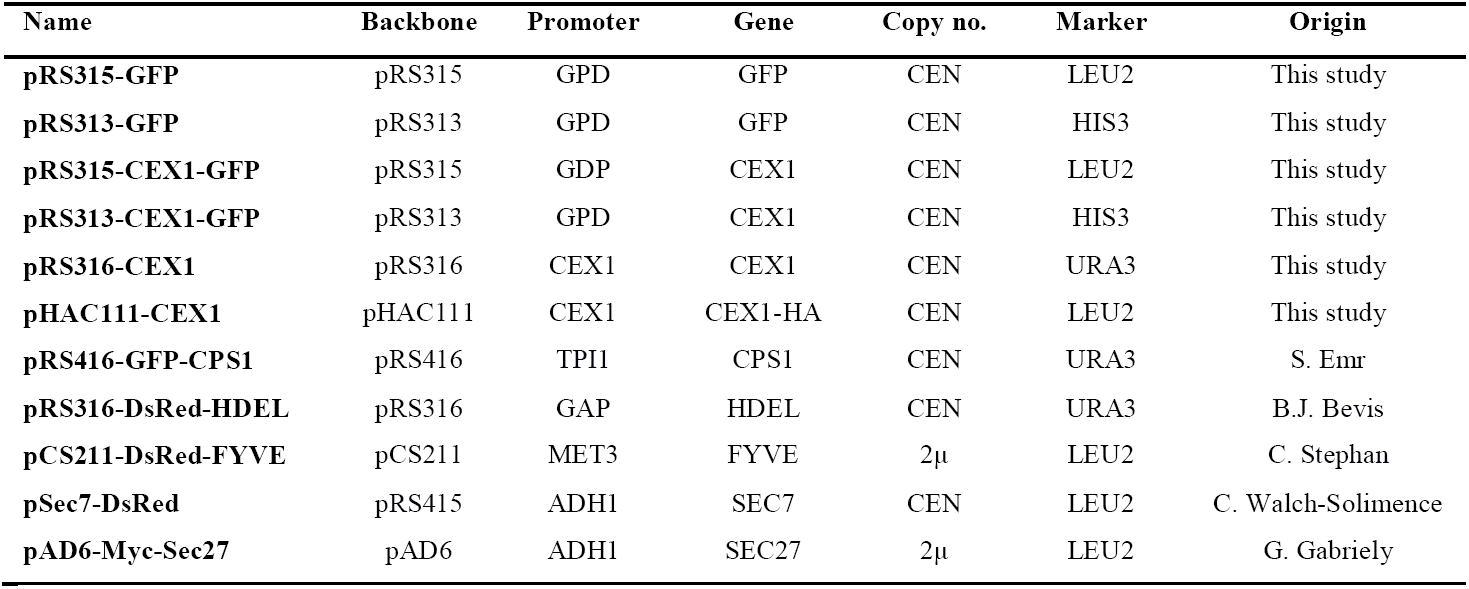
Plasmids used in this study

**Figure 2:**
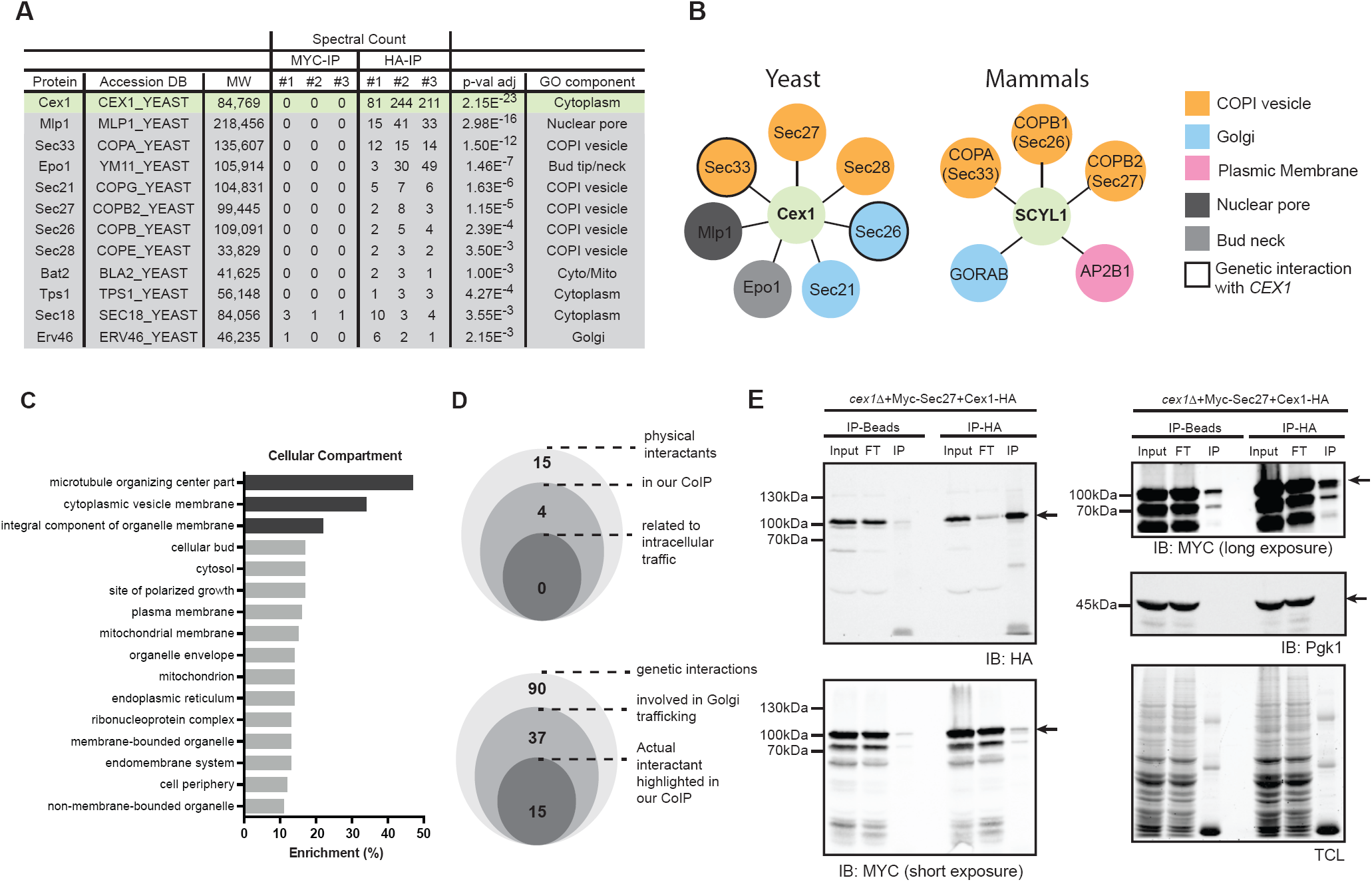
Cex1 interactome reveals its interaction with COPI coat proteins and participation to intracellular trafficking. **A** Summary Table displaying 11 interesting partners identified by CoIP using a *cex1*Δ strain expressing the fusion protein Cex1-HA expressed from its own promoter. **B** Comparison of yeast (Cex1) and mammalian (SCYL1) interactome. Proteins from COPI vesicles are shown in orange, Golgi proteins in blue, proteins from plasmic membrane are in pink and proteins from the nuclear pore or from the bud neck are in grey and light grey respectively. Genes genetically linked to *CEX1* based on the work of Costanzo and colleagues, Science. 2006 are circled. **C** Cellular components analysis of Cex1 interactants in respiration. Enriched GO terms relative to “microtubule organizing center part”, “cytoplasmic vesicle membrane” and “integral component of organelle membrane” were highlighted in grey. **D** Cex1 physical and genetic interactants from the literature (BioGrid and SGD) were compared to our Co-IP data. Fifteen physical partners were identified so far for Cex1, of which 4 were retrieved in our experiments. None of these interactants belongs to intracellular traffic (Upper scheme). Among the 90 genes shown to be genetically linked to *CEX1*, 37 are related to the Golgi and intracellular trafficking, and 15 of them are also Cex1 physical partners based on our experiments (Lower scheme). **E** Co-immunoprecipitation performed with a *cex1*Δ strain expressing the fusion protein Cex1-HA together with a Myc-tagged version of Sec27. Immunoprecipitations were done against HA, and a control IP was performed on non-coated beads. Immunoblotting against HA and MYC revealed respectively the presence of Cex1 and Sec27. Anti-Pgk1 was used as control to assess non-specific adsorption on HA or MYC-coated beads. Total protein loading for each lane was revealed by the stain free technology. Arrows indicate the protein of interest.

A co-IP assay was done to validate the interaction between Cex1 and the Sec27 COPI subunit (Fig 2F). Protein lysates of *cex1*Δ+pHAC-CEX1+pAD6-MYC-SEC27 cells were incubated with Sepharose beads coated with anti-HA or without antibodies (IP-Beads control), and the immunoprecipitates were analyzed by western-blot with anti-HA or anti-MYC antibodies (Fig 2F). In these experiments, the cytosolic protein Pgk1 was used as negative control. These different results show that Cex1 interacts with the COPI coat complex and might be involved in membrane biogenesis and intracellular trafficking.

### Cex1 is associated to membrane fractions

The nature of the association between Cex1 and components of COPI-coated vesicles has to be elucidated. To better understand its link to these vesicles, we performed subcellular fractionation on different yeast strains and followed Cex1-HA distribution (pHAC111-CEX1) by anti-HA western-blot (Fig 3). Subcellular fractionation was performed by differential centrifugation yielding 4 fractions, the total S5 (supernatant 500 ×*g*), two membrane: P13 and P100 (pellet 13000 ×*g* and 100000 ×*g*) fractions and the cytosolic S100 fraction (supernatant 100000 ×*g*). As expected, the cytosolic marker Pgk1 was recovered in the S100 soluble fraction, and the Golgi marker Emp47 in the P13 and P100 membrane fractions (Fig 3A, 3B). In the *cex1*Δ and wild-type (SEY6210) yeast cells, Cex1-HA was mainly detected in the P13 membrane and in the S100 cytosolic fractions, and to a lesser extent in the P100 fractions (Fig 3A). To determine whether this membrane association of Cex1 was mediated by its interaction with the COPI coat complex, we analyzed the subcellular fractionation in strains mutated or deleted in genes encoding for COPI subunits interacting with Cex1 (*sec27-1, sec28*Δ or *sec33-1*). Since the endosomal ESCRT-0 (Endosomal Sorting Complex Required for Transport) subunit Vps27 also interacts with the COPIb subunits Sec27, Sec28 and Sec33 (Gabriely *et al*, 2007), we also tested the *vps27*Δ strain. Compared to the wild-type (SEY6210) and *cex1*Δ strain, the level of soluble Cex1-HA was decreased in the S100 cytosolic fraction of the *sec27-1* and *sec33-1* mutant cells, and to some extent in the *vps27*Δ cells. These data show that the P13 membrane association of Cex1 does not rely on its interaction with COPI subunits. Interestingly, the main phenotype of these COPI mutant strains is the accumulation in the P13 membrane fractions of protein cargos directed to the vacuole, like the carboxypeptidase CPY for example (Gaynor et al., 1997). The overall level of Cex1-HA was significantly reduced in the *sec33-1* mutant (Fig 3A). Based on the spectral counts after the IP (Fig 2A), among the different COPI components Cex1 interacts mainly with Sec33 (Fig 2B), and the loss of Sec33 in the *sec33-1* mutant cells might have an effect on Cex1-HA levels by affecting its stability or turnover.

**Figure 3:**
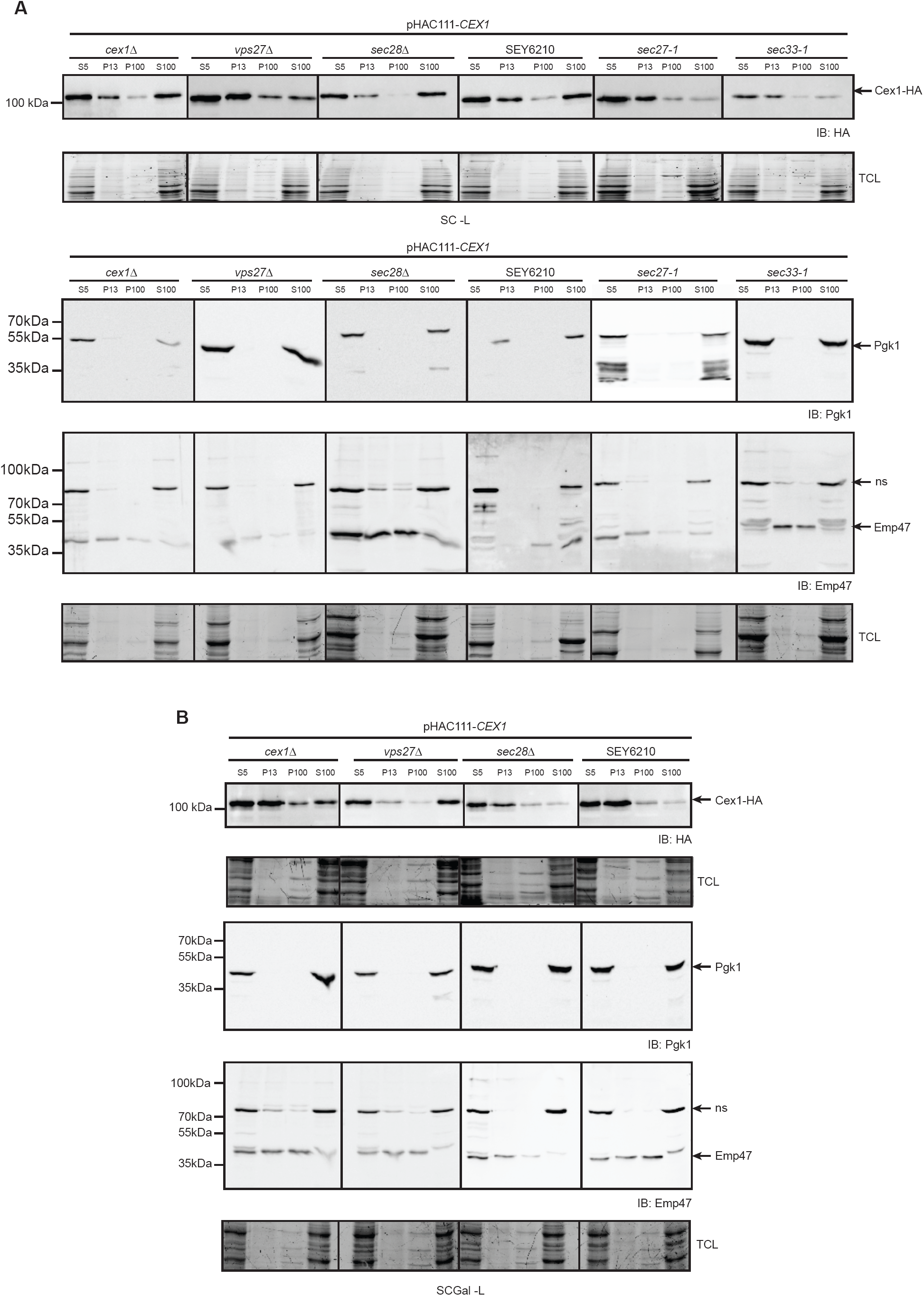
Cex1 is associated to membrane fractions. **A-B** Subcellular fractionation of the WT SEY6210 strain, the control *cex1*Δ strain, compared to the *vps27*Δ, *sec27-1, sec28*Δ and *sec33-1* strains expressing the fusion protein Cex1-HA, and grown in fermentation (SC-Glc; **A**) or in in semi-respiratory medium (SC-Gal; **B**). Localization of Cex1 in each fraction was verified by immunodetection using anti-HA antibodies. Protein loading of each fraction was controlled by stain free gel. Contamination by cytosolic proteins was assessed by detecting the soluble Pgk1 protein, and presence of membrane fraction was evaluated by detection of the ER-to-Golgi and endosomal protein Emp47. ns: non-specific.

Since the expression level of *CEX1* is higher is respiratory conditions (Fig 1D), we also determined the distribution of Cex1 in these conditions. The *sec27-1* and *sec33-1* mutant cells were not able to grow under respiratory conditions (SC-Gly) or under fermentation conditions in which mitochondria are not repressed (SC-Gal) (Fig EV2). This defect was previously reported and is associated to their role in the intracellular localization of mRNAs encoding mitochondrial proteins being imported into mitochondria (Zabezhinsky et al., 2016). We therefore conducted the subcellular fractionation in the Galactose fermentation medium that allows mitochondrial metabolism with the *cex1*Δ, SEY6210, *vps27*Δ and *sec28*Δ strains that grew well in the SC-Gal medium (Fig EV3). In these conditions, Cex1 was detected in the P13 membrane for all strains except for the *vps27*Δ mutant (Figure 3B), suggesting that in these conditions the endosomal Vps27 protein interacting with COPIb subunits might play a role in associating Cex1 to the membrane, however we did not find Vps27 in our interactome study done in respiratory conditions. Interestingly, Cex1 was almost absent from the S100 cytosolic fraction in the SEY6210 and *sec28*Δ strains compared to the *cex1*Δ strain or to the SEY6210 and *sec28*Δ strains grown in fermentation conditions (Fig 3). These data suggest that when mitochondria are not repressed, Cex1 role is mainly associated to the P13 membrane fraction in which the COPI components are also present. All together, these subcellular fractionation data show that COPI complexes integrity governs accurate Cex1 distribution between the membrane and cytoplasmic fractions.

### Intracellular localization of Cex1

Cex1 is interacting with subunits of the COPI coat that are required for ER-to-Golgi retrograde vesicle trafficking. Given its higher expression in respiration, we first analyzed the impact of Cex1 on mitochondria, and showed that the absence of *CEX1* does not impact the mitochondrial network shape and transmission to the daughter cells (Fig EV3A). To determine the intracellular localization of Cex1, we constructed a C-terminally tagged Cex1-GFP fusion protein since we expected that, based on the tridimensional 3D-structure of Cex1, linking GFP after the disordered C-terminal domain of Cex1 would not impair its cellular functions (Fig 4A; Nozawa *et al*, 2013). Next, we analyzed Cex1 cellular localization in fermentation and respiration conditions using the Cex1-GFP fusion protein expressed in the *cex1*Δ strain (See Table 1 and 3, Fig 4B). In these conditions Cex1-GFP was mainly localized in punctuated structures in fermentation and in the cytoplasm and as punctuate structures under respiratory conditions. We sought to determine if the endogenous localization of Cex1 was similar to that of the overexpressed version. To do so, we used a strain in which *CEX1* was genomically GFP-tagged. By confocal microscopy followed by Z-stack acquisition and 3D reconstruction, the presence of Cex1-GFP in puncta was confirmed (Fig 4C, left panels). However the very low GFP signal, most probably due to the low abundance of Cex1 molecules in cells (Ghaemmaghami *et al*, 2003), together with the presence of a strong cytoplasmic GFP signal in respiration, made this analysis less accurate. Nevertheless, merging Z-stack projections of the Cex1-GFP signal clearly showed that in both conditions, Cex1 forms punctate structures in the cytoplasm of yeast cells (Fig 4C, right panels), suggesting that Cex1 could be localized to COPI vesicles.

**Figure 4:**
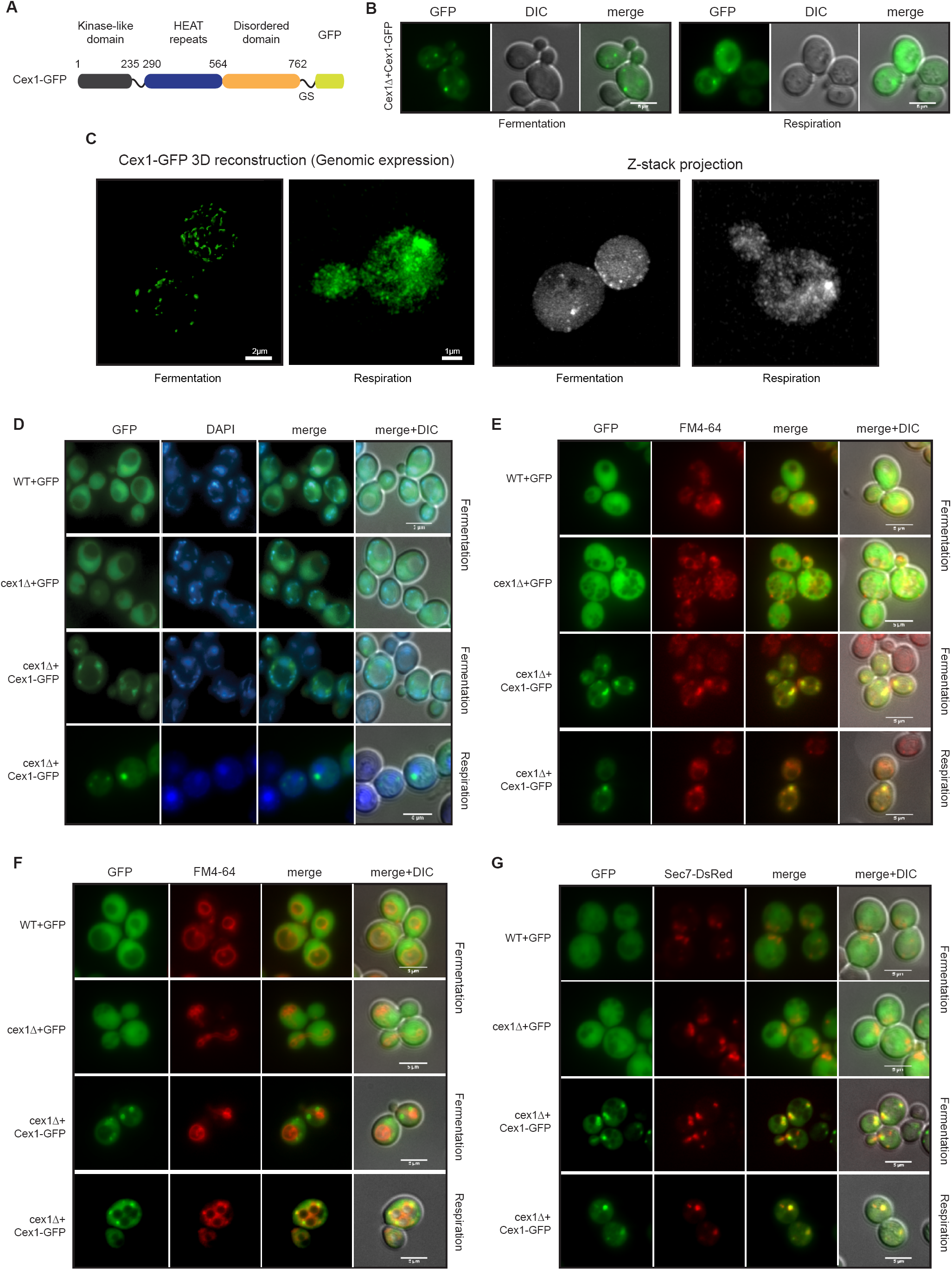
Microscopy-based localization pinpoints Cex1 to the *trans-*Golgi network and in Golgi-to-vacuole endosomes. **A** Scheme of the construct designed to establish Cex1 localization. Expression is controlled by the PGK (phosphoglycerate kinase) constitutive promoter, and Cex1 is linked to the GFP by a glycine-serine (GS) domain. **B** Epifluorescence microscopy of Cex1-GFP expressed in a *cex1*Δ strain. Cells were grown in fermentation and respiration (left and right panels respectively). **C** 3D reconstitution of the Cex1-GFP construct when expressed at its genomic locus (left panels). Images were taken in fermentation and respiration. Corresponding raw images were assembled by Z-stack and each stack fluorescent signal was cumulated by mean intensity (right panels). **D-G** Subcellular localization of Cex1-GFP was performed by looking at 4 compartments. Nuclei were highlighted by DAPI staining (**D**), endosomes (**E**) and vacuoles (**F**) were stained by FM4-64, and the trans-Golgi network and endosomes were visualized by expressing Sec7 fused to Ds-RED (**G**). WT and *cex1*Δ cells expressing GFP alone were used as control. Cells were grown in fermentation and respiration in every condition. Scale bar: 5 µm

We then tracked the subcellular localization of Cex1 by using the Cex1-GFP fusion protein expressed in the *cex1*Δ strain. DAPI staining done on living yeast cells expressing Cex1-GFP or GFP (as a negative control) showed that the fluorescent dots that correspond to Cex1-GFP, colocalized neither with nuclei (round fluorescent signal) nor with mitochondria (Fig 4D). The COPI vesicles are required for Golgi to ER retrograde trafficking of proteins having the HDEL retrieval motif. In wild-type (WT) and *cex1*Δ mutant cells, the HDEL-DsRed is localized at the ER (Fig EV3B), showing that Cex1 is not required for COPI-dependent HDEL retrieval. Moreover, the Cex1-GFP signal did not colocalize with HDEL-DsRed (Fig EV3B). These results show that Cex1 does not belong to the COPI coat required for ER to Golgi transport. The Sec27, Sec28 and Sec33 subunits form the COPIb subcomplex, shown to be involved in post-Golgi trafficking and endosomal sorting (Gabriely *et al*, 2007). Interestingly, we have shown that Sec27, Sec28 and Sec33 interact with Cex1 (Fig 2). Endosomes and vacuoles were stained with the FM4-64 lipidic dye that is internalized by endocytosis (Vida & Emr, 1995). Two incubation times were used in order to stain endosomes (Fig 4E) or vacuoles (Fig 4F). We observed that Cex1 co-localizes with endosomes stained with FM4-64 but not with the vacuolar membrane. To determine whether Cex1 is also localized at the *trans*-Golgi network (TGN) as the COPIb subcomplex (Gabriely *et al*, 2007), we co-transformed the yeast cells with the Sec7-DsRed TGN-localized yeast protein (Richardson *et al*, 2016). Our microscopy observations show that Cex1 partially co-localized with the Sec7-DsRed punctuated structures present in the cells, both in fermentation and respiration (Fig 4G). These results suggest that Cex1 could interact with COPI proteins at the TGN and play a role in protein trafficking at the TGN-endosomal level in yeast cells.

### The Golgi-to-vacuole trafficking pathway is impaired in the absence of *CEX1*

Most vacuolar hydrolases are transported from the Golgi apparatus to the vacuoles *via* the endosomes. The carboxypeptidase Y (CPY) is a vacuolar serine protease, synthesized as a large inactive precursor, which undergoes specific maturation at the ER (p1 form of CPY) and the Golgi (p2 form) apparatus of the cell prior to being fully maturated by cleavage to produce the mature (mCPY) form into the vacuolar lumen. Deterioration of this process leads to several defects among them secretion of unprocessed CPY into the extracellular medium (Stevens *et al*, 1982; Johnson *et al*, 1987). To test whether Cex1 participates to the Golgi-to-vacuole trafficking, we performed a CPY secretion test on solid YPD and YPGly media (Fig 5A, Figure EV4A). In fermentation, the *cex1*Δ strain exhibited a significant higher amount of CPY secretion in contrast to the WT strain (Fig 5A-B). Interestingly, this secretion phenotype was restored by overexpressing Cex1-GFP (Figure 5A-B). We sought to perform the same analysis in respiratory conditions, however the WT strain displayed a high level of CPY secretion similar to the one observed for the *vps27*Δ strain (Fig EV4A), implying that in this condition CPY sorting is somehow affected as it does not follow the same trend than in fermentation, and therefore could not be analyzed. This general defect was further confirmed by the fact that in respiration, p2CPY was recovered in culture supernatant for the WT strain, in cells lacking the proteinase A (*prA*Δ) known to be involved in CPY maturation (Fig EV4B). In the *cex1*Δ strain, we recovered p2CPY but also mCPY in the culture supernatant (Fig EV4B).

**Figure 5:**
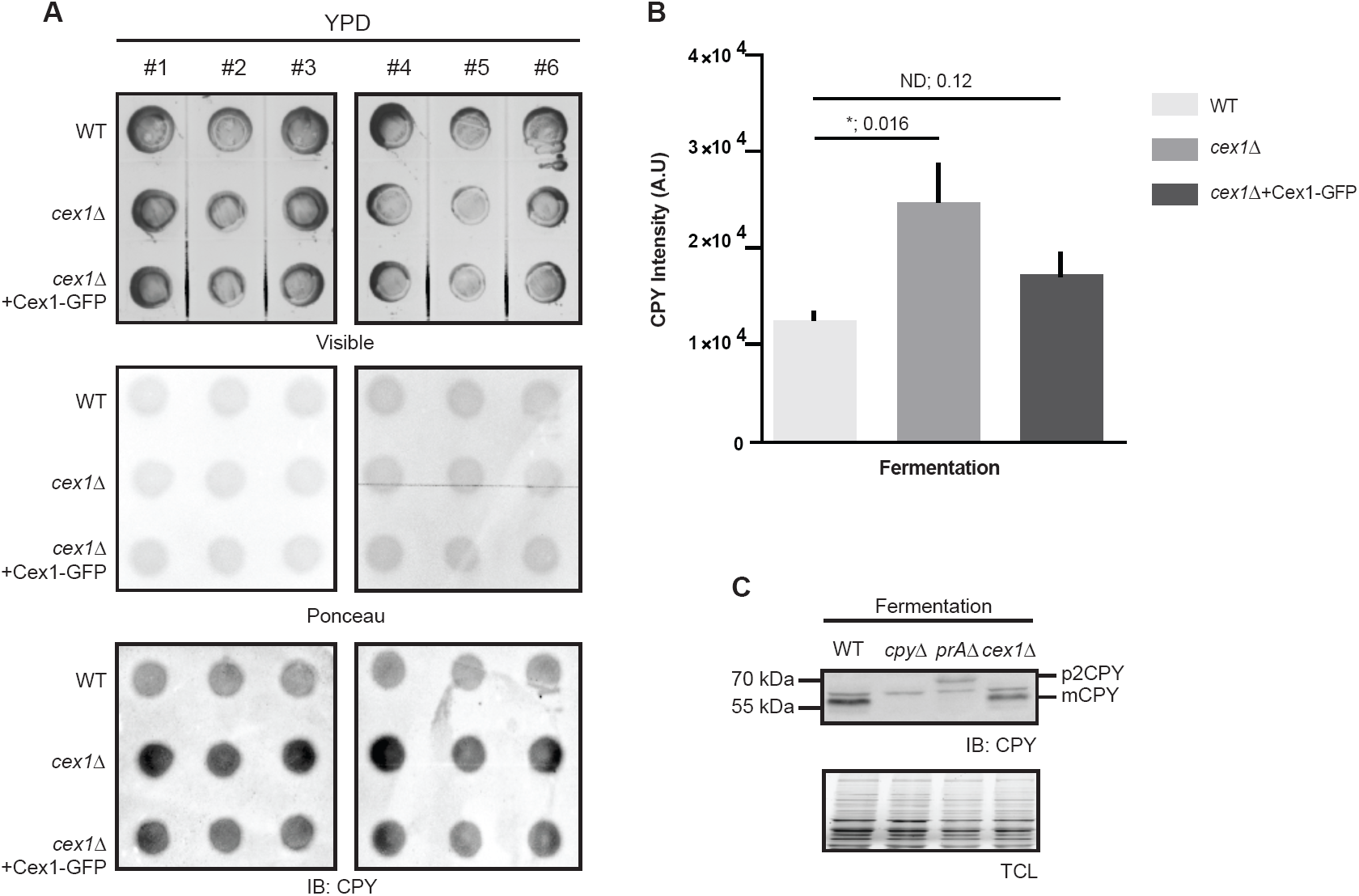
Disruption of *CEX1* leads to CPY extracellular secretion. **A-B** Measurement of CPY secretion in the WT, *cex1*Δ and *cex1Δ*+Cex1-GFP strains when cells were grown in fermentation. Measurements were done based on immunodetection of CPY after solid on-plate secretion assay. **C** CPY maturation assessed by immunodetection on total cell extract under fermentative conditions, in a WT strain, in cells deprived of the gene coding for CPY (*cpy*Δ) or the proteinase A (*pra*Δ), and in the *cex1*Δ strain. Pre-mature (p2CPY) and mature CPY (mCPY) are highlighted. TCL: total cell loading; nd: not different; * *p* val<0.05.

Maturation of the CPY protein was then assessed by following the intracellular level of mCPY in fermentation and respiration conditions. As expected, CPY was efficiently matured in wild-type, absent in *cpy*Δ and not matured in *prA*Δ cells (Fig 5C, Fig EV4C). In these conditions, we did not observe any defect in CPY maturation in the *cex1*Δ strain in respiration (Fig EV4C). However, in fermentation, the ratio between the p2CPY Golgi precursor and the mCPY mature vacuolar was different in *cex1*Δ compared to WT cells (Fig 5C). These data show that in absence of the Cex1 protein, the trafficking of CPY from the Golgi to the vacuole is delayed. We also tested whether in *cex1Δ* cells, the late endosomal or multivesicular body sorting of the carboxypeptidase S (Cps1) was not impaired, our data show that Cps1-GFP is localized to the vacuolar lumen in both WT and *cex1Δ* cells (Fig EV4D). This implies that absence of *CEX1* provokes a delay in the Golgi-to-vacuole trafficking pathway.

Since we could not test the impact of Cex1 on trafficking by following CPY secretion in respiration, we determined Cex1 localization in *vps10*Δ cells. The Vps10 protein is a sorting receptor that shuttles from the TGN to the endosomes and is involved in vesicular trafficking, and cells lacking *VPS10* show mistargeting and secretion of many vacuolar hydrolases (Marcusson *et al*, 1994). We thus asked whether Vps10 influences Cex1 localization in fermentation or respiration by expressing Cex1-GFP in a *vps10Δ* strain. In both conditions, deletion of *VPS10* did not impact the cytosolic content of Cex1 (Fig 6A), but rather increased the number of puncta in cells. Once again, these puncta colocalized with late endosomes in respiration. To confirm this observation, we generated 3D representations of *vps10Δ* cells expressing Cex1-GFP to get rid of the cytosolic signal as well as the background noise induced by classical epifluorescence microscopy. By doing so we unambiguously showed that there were, overall, more puncta in the *vps10Δ* cells than in *cex1Δ* deficient cells, in fermentation (Fig 6B, panels 1-2), but also in respiration (Fig 6B, panels 3-4), suggesting that Cex1’s localization is somehow disturbed in the absence of Vps10. This aberrant localization was further assessed by subcellular fractionation. Consistent with the previous observations, Cex1 was no longer retrieved in the soluble S100 fraction and concomitantly increased in the P100 fraction in which it is almost absent in WT cells (Fig 6C). These data suggest that Vps10 regulates the transport or recycling of Cex1 at the TGN or late endosomes, and lead us to propose that Cex1, through its interaction with Sec27, Sec28, Sec33 and with the help of Vps10, is associated to COPI vesicles to control the sorting of protein cargos.

**Figure 6:**
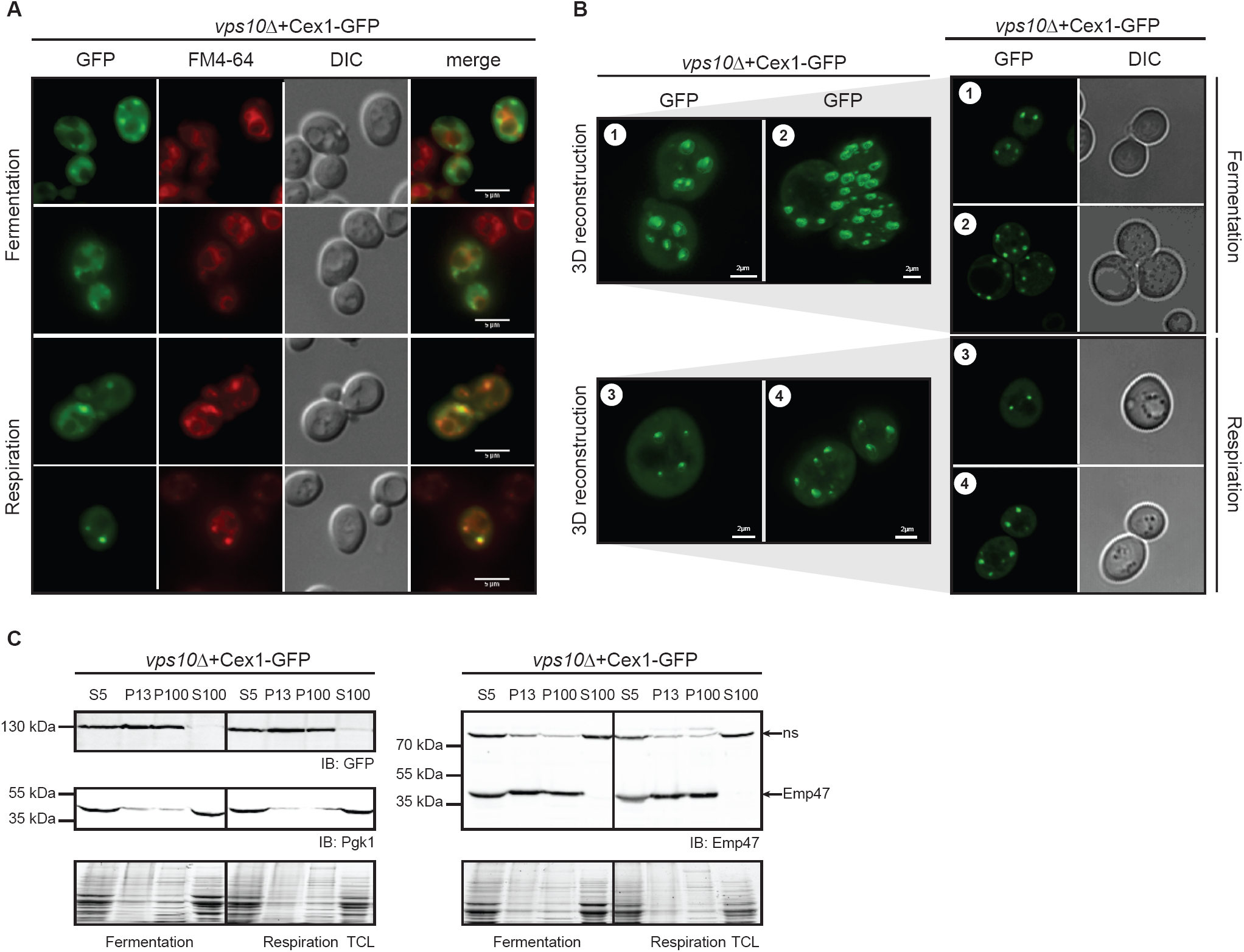
Cex1 endosomal localization is influenced by Vps10. **A** Localization of Cex1-GFP in *VPS10*-deficient cells by epifluorescence microscopy in fermentation and respiration. In both conditions vacuoles and endosomes were stained by FM4-64 for 15 min. **B** Cex1-GFP localization in *vps10Δ* cells visualized after 3D reconstruction of Z-stack images. Two panels were taken in fermentation (1-2) and in respiration (3-4). For each panel (1-2-3-4), one section of the Z-stack was used to show GFP localization and the corresponding DIC image. Scale bar 2 µm. **C** Subcellular fractionation of *vps10Δ* cells expressing Cex1 coupled to GFP. Cex1 sublocalization was followed by immunodetection using anti-GFP antibodies. Anti-Pgk1 and anti-Emp47 antibodies were respectively used as cytosolic and ER/Golgi marker, to ascertained proper fractionation.

## Discussion

Based on previous reports, we sought to identify the genetic link between *ARC1* and *CEX1* in the nuclear export of aa-tRNA (McGuire & Mangroo, 2007; 2012; Nozawa *et al*, 2013). To do so we generated a strain deprived of both *CEX1* and *ARC1* using the same genetic background as previously described (McGuire & Mangroo, 2007). However, as we clearly showed here, there is no genetic link between these genes, at least in the conditions we tested. Besides our report, other teams also demonstrated the absence of a genetic link between *ARC1* and *CEX1* (Costanzo *et al*, 2016). Of note, a recent large-scale genetic interaction network consisting in nearly 1 million genetic interactions was constructed in *S. cerevisiae* (Costanzo *et al*, 2016). This high-throughput study using deletion and thermo-sensitive strains does not report a genetic link between *ARC1* and *CEX1.* This finding challenges the biological role described for Cex1 in the export of aa-tRNA in *S. cerevisiae* and raised several questions. Indeed, since Cex1 is most likely not involved in the nuclear export of aa-tRNA, we analyzed the localization of Cex1, its interaction network and hence its cellular function in yeast. Here we show that Cex1 is a cytoplasmic protein that interacts with components of the COPI vesicular coat and localizes at the TGN and endosomes in fermentation and respiration conditions (Fig 2, 3 and 4). Interestingly in respiration, a higher fraction of Cex1 is retrieved in the cytoplasm. Therefore we can speculate that Cex1 display different roles, more likely accomplished by the same molecular function, when associated to vesicles or as “free” particle in the cytoplasm. Consistent with our present findings, SCYL1, the orthologue of Cex1 in mammals, is also a constituent of vesicular complexes and regulates Golgi-to-ER transport and Golgi morphology both in Human and mice *via* its interaction with COPI coat proteins (Burman *et al*, 2008; 2010). The same role has also been suggested through an ORFeome analysis of GFP-tagged proteins for the orthologue Ppk3 in the fission yeast *S. pombe* (Matsuyama *et al*, 2006).

Based on our data, we envision 2 scenarios for the molecular and cellular function of Cex1. Cex1 could act as hub *via* its HEAT-repeats domain to recruit COPIb proteins at the TGN, or at the level of early endosomes, to ensure Vps10-dependent sorting of vacuolar hydrolases trafficking from the TGN to the vacuole. The second hypothesis is based on the recent observation that pseudokinase domains, despite lacking their catalytic activity, still retained the ability to bind and/or hydrolyze ATP (Murphy *et al*, 2014). This ATP-binding property is sought to regulate a proportion of pseudokinase-dependent signaling, either through modulation of catalytic activity, or by conformational transition (Byrne *et al*, 2017; Hammarén *et al*, 2015). Cex1 could then act as an ATP-exchange factor, as a substrate trap or as a mobile platform to promote protein-protein interactions at the TGN and early endosomes. Regardless of its genuine molecular function, Cex1 regulates the vesicular trafficking and sorting from the TGN to the vacuole. This is supported by several lines of evidence: (*i*) Cex1 co-localizes with the TGN Sec7 protein and with FM4-64-stained endosomes (Fig 4), (*ii*) cells lacking *CEX1* exhibit CPY secretion and accumulation of Golgi p2CPY, two phenotypes characteristic of *vps* (vacuolar protein sorting) mutants (Fig 5), and (*iii*) Cex1 accumulates on membrane structures in a *vps10*Δ strain lacking the receptor required for the transport of the vacuolar hydrolases from the TGN to the endosomes (Fig 5). Among the GO terms associated to cellular compartments, we found terms related to the MTOC (microtubule-organizing center) enriched in our CoIP, notably through to the interaction of Cex1 with Spc42, Spc110, Mps2 and Cnm67 (Table EV1). In *S. cerevisiae*, mitosis takes place inside the nucleus and the centrosome (MTOC) is assembled by two centrioles. During cellular division nuclei dissociate, and microtubules interact with duplicated chromosomes to transport them to the mother and daughter cell along the mitotic spindle. According to Kegg (http://www.genome.jp/kegg/), Cex1 has a paralogue, Pds5 that acts as a cohesion maintenance factor, involved in sister chromatid condensation and cohesion (Hartman *et al*, 2000; Panizza *et al*, 2000; Stead *et al*, 2003). In terms of structure, they both share the Armadillo-type fold retrieved in HEAT repeats of Cex1 (Hartman *et al*, 2000; Nozawa *et al*, 2013). In addition, a thermo-sensitive mutant of Pds5 (*pds5-3*) accumulates chromatin at the bud neck (Hartman *et al*, 2000). This led us to hypothesize that, like for one transcriptional variant of its human homologue *SCYL1* (SCYL1_4; Kato *et al*, 2002), Cex1 might somehow be involved in chromatin binding. Although we did not localized Cex1 in the nucleus in our experiments, we hypothesize that under appropriate conditions, post-translational modifications might be necessary to recruit a portion of Cex1 to the MTOC and participate, to some extent, to the chromatin separation and transfer to daughter cells during budding, but this has to be further investigated.

SCYL1, SCYL2 and SCYL3 are members of the SCY1-like (SCYL) family of mammalian pseudokinase proteins, and several studies suggested that they somehow participate to the regulation of protein trafficking along the secretory pathway (Conner & Schmid, 2005; Düwel & Ungewickell, 2006; Borner *et al*, 2007; Burman *et al*, 2008; 2010; Hamlin *et al*, 2014). Even if SCYL1 was not detected in a recent proteomic analysis of COPI-coated vesicles (Gilchrist *et al*, 2006), it has to be noted that peripheral membrane proteins associating with organelles are not always identified by MS analysis (Blondeau *et al*, 2004). SCYL1 possess the RKLD-COO-motif, a variant of the dilysine motif KKxx-COO-responsible for COPI binding (Burman *et al*, 2008). It has also been shown that internal KK motifs present in the C-terminal helical region are involved in the binding of cytoplasmic proteins to β’COP protein (Sullivan *et al*, 2000). Although Cex1 lacks one of the COPI-binding motifs in its C-terminal part, its helical region is rich in KK motifs, which further rationalizes the observation that Cex1 is interacting with COPI vesicles.

Development of neurodegenerative diseases is sometimes associated with dysfunction of the intracellular trafficking apparatus (Ferrucci *et al*, 2011; Majcher *et al*, 2015; Dahm & Macchi, 2012), and in human as well as in mouse mutations in the *SCYL1* gene have been linked to amyotrophic lateral sclerosis (ALS) (Pelletier *et al*, 2012; Schmidt *et al*, 2015). In human *SCYL1*, three mutations in three individuals from two different families suffering from peripheral neuropathy, cerebellar atrophy, and ataxia were identified (Schmidt *et al*, 2015). One frame-shift mutation identified in mice *SCYL1* leads to a loss-of-function and this mutation is called muscle deficient (*mdf*), defined by an early onset of progressive motor neuron disorder (Schmidt et al., 2007). The *Scyl1*^*mdf/mdf*^ mice display cytosolic accumulation of TDP-43 and Ubiquilin 2 and have similar symptoms to that observed in ALS patients (Pelletier *et al*, 2012). The fact that a protein regulating COPI trafficking and Golgi morphogenesis is somehow involved in ALS both in Human and mice opens a new road to better understand the development of this disease. Unfortunately, there are no studies aiming to dissect the link between SCYL1 intracellular trafficking and ALS.

*SCYL1* is well conserved in eukaryotes and ubiquitously expressed in human cells. It would then be interesting to study the role of SCYL1 or its yeast homologue in the case of ALS in the light of our new findings that describe a role for Cex1 in protein sorting at the *trans*-Golgi-network and the endosomes.

## Materials and Methods

### Media and growth condition

The following rich media were used for the growth of yeast strains: 1 % (w/v) yeast extract, 1 % (w/v) peptone, 40 mg/L adenine, 2 % (w/v) glucose (Glc), 2 % (w/v) or 2 % (w/v) glycerol (Gly). We also used complete synthetic medium (SC) composed of 0.17 % (w/v) yeast nitrogen base without amino acids and ammonium sulfate, 0.5 % (w/v) ammonium sulfate, 2% (w/v) Glc and 0.8 % (w/v) of a mixture of amino acids and bases from Formedium (Norfolk, UK). The solid media contained 2 % (w/v) agar. Every strains were grown at 30°C with rotational shaking to mid-log (OD_600nm_ 0,7) or log phase (OD_600nm_ 1.5) depending on the experiments. Prior to subcellular fractionation, thermo-sensitive strains (*sec27-1* and *sec33-1*) were grown at 25°C to mid-log phase, and then transferred at 37°C for 3h.

### Drop tests and CPY secretion assay

Drop tests were done using 10 mL cultures grown to log phase. Cells were then spun down, diluted in water to a final OD_600nm_ 0.5 and further diluted to the tenth 4 times. 7 μL of each dilution was spotted onto agarose plates and incubated for at least 2 days at 25, 30 or 37°C.

For secretion test cells were spotted onto YPD and YPGly agarose plates similarly to a drop test. Once dry, cells were covered with PVDF membrane and incubated for 2 to 4 days at 30 °C. PVDF membranes were then washed once with water, and blocked with 5 % milk in TBS-Tween buffer, and presence of the CPY was assessed by Western-blot using a rabbit anti-cpy antibody (1:5,000; Santa Cruz), and HRP-conjugated anti-rabbit antibodies (1:10,000; BioRad).

### RNA extraction and qPCR experiments

All total mRNA samples were prepared with trizol^®^ (Invitrogen), all cDNAs were prepared using AMV reverse transcriptase (MP biomedicals) with 6 μg of total RNA and a 16-mer poly-dT oligoDNA, following recommendations of manufacturers. All q-PCR reaction mixtures were performed with 5 × HOT Pol^®^ EvaGreen^®^ qPCR Mix Plus Rox (Euromedex) and quantifications were performed in a MX3005P thermocycler (Stratagene). Data were analyzed with the REST software (Pfaffl et al., 2002). *ACT1, NUP57* and *NOP1* were used as internal references. Measurements of *ARC1* and *CEX1* mRNA transcripts in various knock-out strains were also performed on 50 mL cultures grown in fermentation (SC-Glc) or respiration (SC-Gly).

### Yeast transformation

Fifty mL of yeast cells were grown in SC-Glc to mid-log phase at 30°C. Cells were spun down and washed in 1 volume of deionized water (5 min at 3000 *×g* at room temperature). The pellet was then resuspended in 5 mL of lithium acetate mix (1× TE; 100 mM lithium acetate), spun down (5 min at 3000 *×g* at room temperature) and the pellet was resuspended in 0,5 mL of lithium acetate mix and transferred to an Eppendorf tube. After 1 hour of shaking (orbital, 150 rpm, 30 °C), 200 μL of cells were mixed with 10 μL of 10 mg/mL boiled sheared salmon sperm DNA and 1-5 μg of DNA (PCR product or plasmid). The mix was incubated for 30 min on an orbital shaker (150 rpm, 30°C), 1 mL of PEG3500 was added and the mix was again incubated for 30 min. Following this, cells were incubated 10 min at 42°C, spun down (5 min at 3000 *×g* at room temperature), pellet was resuspended in 200 μL of TE buffer (100 mM Tris-HCl pH 7.5; 10 mM EDTA pH 8) and cells were plated onto selective media and incubated at 25 or 30°C depending on the strain.

### *ARC1* deletion in the Δ*cex1* strain

The *cex1*Δ strain (see Table 1) was used to obtain the *cex1*Δ *arc1*Δ. A *HIS3* cassette containing *S. cerevisiae HIS3* gene with its promoter (312 bp) and terminator (201 bp) was generated by PCR. Using the ARC1-HIS3 Fwd and ARC1-HIS3 Rev primers (See table 2), 50 bp of homology domain from *ARC1* localized at the ATG and STOP codons were appended. The *HIS3* cassette was amplified using 1 μL of *S. cerevisiae* genomic DNA (50 ng/μL), 0.3 μM of each primer, 4 μL of Phusion Buffer (10×), 0.2 μL of dNTP (25 mM) and 1 U of Phusion DNA polymerase (Thermo Scientific) in a 20 μL reaction mix. Reaction was performed 30 sec at 98°C, then 10 sec at 98°C, 20 sec at 60°C and 30 sec at 72°C for 30 cycles, and ended by 3 min of reaction at 72°C. After PCR purification using the Wizard^®^ SV Gel and PCR Clean-Up System (Promega), 1 μg of PCR product was then transformed in *cex1*Δ +pRS316-CEX1 strain.

### Proteins extraction and western blots

Fifty (50) mL of cells grown to log phase were harvested, and resuspended in 2 mL of breaking buffer (100 mM Tris-HCl pH6.8, 150 mM NaCl, 0.5 mM EDTA), additioned with a cocktail of anti-proteases (cOmplete Mini, EDTA-free – ROCHE) and glass-beads. Cells were broken by mechanical disruption (6 cycles at 6 m/sec during 30 sec) using Fast-Prep (MP Bio). Each aliquot was then incubated 30 min on ice in the presence of 0.5 % (v/v) NP40. Protein extracts were obtained after 1 h of centrifugation at 105000 *×g* at 4°C.

Prior to Western-blots, protein concentrations were determined using Bradford, and 10–15 µg of proteins were separated by SDS-PAGE on a 10 % or 12 % gel prior to electroblotting onto Hybond-P membrane (Amersham). Detection was carried out using HRP-conjugated anti-rabbit, anti-mouse or anti-goat antibodies (Bio-Rad), at a concentration of 1:5000. We used ECL-plus reagents (BioRad) according to the manufacturer’s instructions. We used mouse anti-GFP primary antibodies (1:10000) to show the presence of Cex1 in extracts. Presence of Pgk1 was probed with a mouse monoclonal anti-Pgk1 antibody (1:5000; Abcam), Cex1-HA proteins were identified using a mouse monoclonal anti-HA antibody (1:10000). In subcellular fractionation, Emp47 was detected using rabbit polyclonal anti-Emp47 (1:5000; gift from R. Duden).

### Subcellular fractionation

One hundred mL of cells were grown to mid-log phase in appropriate media and harvested by centrifugation 4 min at 1600 *×g* at room temperature. Cells were then washed once in 10 mL of cold lysis buffer (20 mM Hepes KOH pH 6,8; 150 mM KoAc; 10 mM MgCl_2_; 250 mM Sorbitol; protease inhibitor cocktail), resuspended in 1 mL of cold lysis buffer and transferred in tubes containing 1/3 volume of glass beads. Cells were broken by mechanical disruption (6 cycles at 6 m/sec during 30 sec) using Fast-Prep (MP Bio), transferred to a new 1.5 mL Eppendorf tube and centrifuged at 300 ×*g* for 5 min at 4°C. Supernatants were transferred to newly tubes and centrifuged at 500 ×*g* for 5 min at 4°C, pellets were resuspended in 200 μL of lysis buffer (P5), aliquots (100 μL) of supernatants were kept for further analysis (S5) and the remaining supernatants were again centrifuged at 13000 ×*g* for 10 min at 4°C. Pellets (P13) were resuspended in 100 μL of lysis buffer while supernatants were centrifuged for an hour at 100000 ×*g* at 4°C. Supernatants (S100) were kept for further analysis and pellets (P100) were resuspended in 50 μL of lysis buffer.

### Fluorescent microscopy analyses and images acquisition

Cells were incubated overnight in appropriate media, and living cells in exponential phase of growth were used for microscopy studies. For late endosomes and vacuoles staining, cell cultures were centrifuged 1 min at 1500 ×*g* at room temperature, resuspended in 50 μL of YPD encompassing 4 μL of FM4-64 (200 μM, Life Technologies), and incubated 5 to 20 min at rotational shaking at 30°C. Cells were washed in 500 μL of fresh YPD, resuspended in 100 μL of SC-Glc and incubated for another 15 min at 30°C. For vacuolar staining in respiration, cells were treated as described but were incubated 30 min at 30°C, washed in YPGly, resuspended in 100 μL of SC-Gly and incubated for another hour at 30°C. Mitochondria were stained using MitoTracker Red CMXRos (Invitrogen). One μL (200 μM stock) was added to 1 mL of culture and cells were incubated 20 min at 30°C with rotational shaking. Cells were then washed with appropriate medium and images were taken. Nuclei were stained with 15 µL of DAPI (10 µg/mL) and incubated 10 min in the dark. Three washes with PBS are needed to avoid non-specific background fluorescence. Observation was performed with a 100X/1.45 oil objective (Carl Zeiss) on a fluorescence Axio Observer D1 microscope (Carl Zeiss) using DAPI, GPF or DsRED filters and DIC optics. Images were captured with a CoolSnap HQ2 photometrix camera (Roper Scientific) and treated by ImageJ (Rasband W.S., ImageJ, U. S. National Institutes of Health, Bethesda, Maryland, USA, http://imagej.nih.gov/ij/). Images for 3D reconstruction were taken using a confocal LSM 780 high resolution module Airyscan with a 63X 1.4NA plan apochromatic objective (Carl Zeiss) controlled by the Zen Black 2.3 software (Carl Zeiss). Z-stack reconstruction was performed on the IMARIS 9.1.2 (Bitplane AG) software.

### Co-immunoprecipitation

To identify Cex1 partners, 3 independent clones were grown to log phase (1.2) in 50 mL of SC-Gly -L liquid medium and cells were harvested by centrifugation at 4100 *×g* for 10 min at room temperature. Cells were frozen in liquid nitrogen and lysed with a mortar at 4°C. The cell powder obtained was then resuspended in a lysis buffer (0.33 % NP40; 50 mM Tris-HCl pH8; 50 mM NaCl; 1 mM PMSF, supplemented with cOmplete Roche antiproteases, and centrifuged at 12000 ×*g* for 15 min at 4°C. The supernatant was recovered and 1.35 mg of proteins were mixed with 50 μL of magnetic anti-HA MicroBeads (Myltenyi Biotec) and incubated 30 min on ice. The suspension was then loaded onto μMacs column and elution was done following manufacturer’s instructions. The washing step was done using a modified version of the lysis buffer containing 0.1 % of NP40. Final elution was done with 50 μL of Laemli buffer boiled at 95°C.

### Mass spectrometry and Nano-LC/MS analysis

For nano-LC-MS/MS analysis, the dried extracted peptides were transferred in vials compatible with nano-LC-MS/MS analysis (Ultimate 3000, Dionex and MicroTOF-Q, Bruker). The method consisted in a 60-min gradient at a flow rate of 300 nL/min using a gradient from two solvents: A (0.1% formic acid in water) and B (0.08% formic acid in acetonitrile). The system includes: a 300 µm × 5 mm PepMap C18 precolumn (Dionex) in order to pre-concentrate peptides and a 75 µm × 150mm C18 column (Dionex) used for peptide elution. MS and MS/MS data were acquired in a data-dependent mode using Hystar (Bruker) and processed using Mascot software (Matrix Science). Consecutive searches against the NCBI nr database first and then against the *S. cereviaiae* taxonomy were performed for each sample using an intranet version of Mascot 2.0. Peptide modifications allowed during the search were: N-acetyl (protein), carbamidomethylation (C) and oxidation (M). The other parameters were: peptide tolerance = 0.4 Da, MS/MS tolerance = 0.4 Da, 2 missed cleavage sites by trypsin allowed. Proteins showing two peptides with a score higher than the query threshold (p-value<0.05) were automatically validated with Proteinscape (Bruker). Each protein identified by only one peptide was checked manually using the classical fragmentation rules. Protein bands were manually excised from the gels and transferred into 96-well microtitration plates. Excised gel samples were cut in small pieces and washed three times by incubation in 25 mM NH_4_HCO_3_ for 15 min and then in 50 % (v/v) acetonitrile containing 25 mM NH_4_HCO_3_ for 15 min. Samples were then dehydrated with 100% acetonitrile and then reduced with 10 mM DTT during 1 h before being alkylated with 55 mM iodoacetamide for 1 h in the dark. Gel pieces were washed again with the destaining solutions described above. 0.250 µg of modified trypsin (Promega, sequencing grade) in 25 mM NH_4_HCO_3_ were added to the dehydrated gel spots depending on protein amount. After 30 min incubation at room temperature, 20 µL of 25 mM NH_4_HCO_3_ were added on gel pieces before incubation overnight at 37°C. Peptides were then extracted from gel pieces in 20 µL of 50 % acetonitrile / 5 % formic acid.

### Statistics

Statistical analyses were done with GraphPad Prism 6 using the unpaired t-test. Every experiment was done in biological triplicates, and SEM are showed on every graph.

## Acknowledgements

We thank Pr. J. Gerst for sharing the pAD6-MYC-SEC27 plasmid and Pr. A. Spang for the *sec27-1* and *sec33-1* strains and CPY antibodies. We also thank Pr. R. Duden for providing Emp47 antibodies and Pr. C. Ungermann for providing the *vps10Δ* strain. We thank Dr. Robert Martin and Frederic Fisher for insightful discussions. We thank E. Vega (Plateau d’imagerie cellulaire I2MC Toulouse INSERM UMR1048 – TRI Génotoul) for technical help on Airyscan images acquisition and 3D reconstruction, and Dr. R. Gaudin (Institute of viral and liver disease, UMR-S1110, Strasbourg) for providing kind access to IMARIS software. This work was supported by the French National Program Investissement d’Avenir administered by the ‘‘Agence National de la Recherche’’ (ANR), ‘‘MitoCross’’ Laboratory of Excellence (Labex), funded as ANR-10-IDEX-0002-02 (to H.D.B, L.E, B.S, Y.O.C), the Agence Nationale de la Recherche ANR-13-BSV2-0004 (to S.F), the JST-CNRS Japanese-French Cooperative Program on “Structure and Function of Biomolecules” (O.N. and H.D.B), the University of Strasbourg (H.D.B, L.E, B.S, Y.O.C, P.H, S.F, B.R), the CNRS (H.D.B, L.E, B.S, Y.O.C, P.H, S.F, B.R). The mass spectrometry instrumentation was funded by the University of Strasbourg, IdEx “Equipement mi-lourd” 2015.

## Author contributions

All authors have seen and approved the manuscript and its contents. LE, JOD, SF, HDB designed the experiments; LE, BR, SF, PH performed the experiments. LE, SF, ON, HDB wrote the manuscript.

## Expanded View Figure legends

**Figure EV1 related to Figure 1: *ARC1* and *CEX1* are not synthetic lethal.**

**A** WT, *cex1*Δ, *arc1*Δ, *cex1*Δ*arc1*Δ, *cex1*Δ expressing Cex1-GFP and *vps27*Δ strains were spotted on fermentative (SC-Glc), respiratory (SC-Gly), with sucrose as the sole source of carbon (SC-Suc) or on medium eliciting osmotic shock (1 M sorbitol) in fermentation and respiration.

**B** Schematic of tRNA species interacting with Arc1 as described in Deinert et al., 2001, and amino acids species stored in the vacuole. In bold are the three species shared by Arc1-bound tRNAs and vacuole storage. WT, *cex1*Δ, *arc1*Δ, *cex1*Δ*arc1*Δ, *cex1*Δ expressing Cex1-GFP and *vps27*Δ strains were spotted on fermentative (SC-Glc) media supplemented with 5 mM of isoleucine (Ile), proline (Pro) or arginine (Arg). All strains were grown at 30°C for 2 days.

**Figure EV2 related to Figure 3: Strains deprived in trafficking-related genes are respiratory deficient.** *Cex1*Δ, SEY6210, *vps27*Δ, *sec27-1, sec28*Δ and *sec33-1* cells were grown in fermentation (SC-Glc), in a semi-respiratory media (SC-Gal) or in respiration (SC-Gly) at 30 and 25°C for 2 to 3 days.

**Figure EV3 related to Figure 4: *CEX1* disruption does not impact other cellular functions than Golgi-to-vacuole trafficking.**

**A** Mitochondria staining by MitoTracker CMXRos in WT, *cex1*Δ and *cex1*Δ+Cex1GFP strains. Cells were grown in respiration.

**B** Subcellular localization of Cex1-GFP by looking at Golgi-to-ER vesicles with the Ds-Red-coupled HDEL retrieval motif. Cells were grown in fermentation and respiration in every condition. Scale bar: 5 µm

**Figure EV4 related to Figure 5: Golgi-to-vacuole trafficking is impaired in *cex1***Δ **cells.**

**A** CPY secretion test performed in respiration (YPGly) after 4 days of growth at 30°C. Three drops of each strain was spotted in duplicate. Cells (visible), total secreted proteins (Ponceau staining) and CPY secretion were monitored.

**B** Immunodetection of CPY in culture supernatant of WT, *cpy*Δ, *prA*Δ and *cex1*Δ strains in respiration. **C** CPY maturation assessed by immunodetection on total cell extract under fermentative conditions, in a WT strain, in cells deprived of the gene coding for CPY (*cpy*Δ) or the proteinase A (*pra*Δ), and in the *cex1*Δ strain. Pre-mature (p2CPY) and mature CPY (mCPY) are highlighted. TCL: total cell lysate.

**D** Localization in fermentation and respiration of the vacuolar carboxypeptidase S (Cps1) fused to the GFP and expressed in WT and *cex1*Δ cells. Scale bar: 5 µm

## References

Audhya A, Foti M & Emr SD (2000) Distinct roles for the yeast phosphatidylinositol 4-kinases, Stt4p and Pik1p, in secretion, cell growth, and organelle membrane dynamics. Mol. Biol. Cell 11: 2673–2689

Bertrand E, Houser-Scott F, Kendall A, Singer RH & Engelke DR (1998) Nucleolar localization of early tRNA processing. Genes & Development 12: 2463–2468

Blondeau F, Ritter B, Allaire PD, Wasiak S, Girard M, Hussain NK, Angers A, Legendre-Guillemin V, Roy L, Boismenu D, Kearney RE, Bell AW, Bergeron JJM & McPherson PS (2004) Tandem MS analysis of brain clathrin-coated vesicles reveals their critical involvement in synaptic vesicle recycling. Proceedings of the National Academy of Sciences 101: 3833–3838

Borner GHH, Rana AA, Forster R, Harbour M, Smith JC & Robinson MS (2007) CVAK104 is a Novel Regulator of Clathrin-mediated SNARE Sorting. Traffic 8: 893–903

Burman JL, Bourbonniere L, Philie J, Stroh T, Dejgaard SY, Presley JF & McPherson PS (2008) Scyl1, mutated in a recessive form of spinocerebellar neurodegeneration, regulates COPI-mediated retrograde traffic. Journal of Biological Chemistry 283: 22774–22786

Burman JL, Hamlin JNR & McPherson PS (2010) Scyl1 Regulates Golgi Morphology. PLoS ONE 5: e9537–15

Byrne DP, Foulkes DM & Eyers PA (2017) Pseudokinases: update on their functions and evaluation as new drug targets. Future Medicinal Chemistry 9: 245–265

Chafe SC, Pierce JB, Eswara MBK, McGuire AT & Mangroo D (2011) Nutrient stress does not cause retrograde transport of cytoplasmic tRNA to the nucleus in evolutionarily diverse organisms. Mol. Biol. Cell 22: 1091–1103

Conner SD & Schmid SL (2005) CVAK104 Is a Novel Poly-l-lysine-stimulated Kinase That Targets the β2-Subunit of AP2. Journal of Biological Chemistry 280: 21539–21544

Costanzo M, VanderSluis B, Koch EN, Baryshnikova A, Pons C, Tan G, Wang W, Usaj M, Hanchard J, Lee SD, Pelechano V, Styles EB, Billmann M, van Leeuwen J, van Dyk N, Lin Z-Y, Kuzmin E, Nelson J, Piotrowski JS, Srikumar T, et al (2016) A global genetic interaction network maps a wiring diagram of cellular function. Science 353: aaf1420–aaf1420

Dahm R & Macchi P (2012) Human pathologies associated with defective RNA transport and localization in the nervous system. Biology of the Cell 99: 649–661

Deinert K, Fasiolo F, Hurt EC & Simos G (2001) Arc1p organizes the yeast aminoacyl-tRNA synthetase complex and stabilizes its interaction with the cognate tRNAs. Journal of Biological Chemistry 276: 6000–6008

Düwel M & Ungewickell EJ (2006) Clathrin-dependent association of CVAK104 with endosomes and the trans-Golgi network. Mol. Biol. Cell 17: 4513–4525

Ferrucci M, Fulceri F, Toti L, Soldani P, Siciliano G, Paparelli A & Fornai F (2011) Protein clearing pathways in ALS. Arch Ital Biol 149: 121–149

Gabriely G, Kama R & Gerst JE (2006) Involvement of Specific COPI Subunits in Protein Sorting from the Late Endosome to the Vacuole in Yeast. Mol. Cell. Biol. 27: 526–540

Gaynor EC, Emr SD (1997) COPI-independent Anterograde Transport: Cargo-selective ER to Golgi Protein Transport in Yeast COPI Mutants. J. Cell. Biol. 136 (4): 789

Ghaemmaghami S, Huh W-K, Bower K, Howson RW, Belle A, Dephoure N, O’Shea EK & Weissman JS (2003) Global analysis of protein expression in yeast. Nature 425: 737–741

Gilchrist A, Au CE, Hiding J, Bell AW, Fernandez-Rodriguez J, Lesimple S, Nagaya H, Roy L, Gosline SJC, Hallett M, Paiement J, Kearney RE, Nilsson T & Bergeron JJM (2006) Quantitative proteomics analysis of the secretory pathway. Cell 127: 1265–1281

Gillooly DJ, Morrow IC, Lindsay M, Gould R, Bryant NJ, Gaullier JM, Parton RG & Stenmark H (2000) Localization of phosphatidylinositol 3-phosphate in yeast and mammalian cells. The EMBO Journal 19: 4577–4588

Hamlin JNR, Schroeder LK, Fotouhi M, Dokainish H, Ioannou MS, Girard M, Summerfeldt N, Melancon P & McPherson PS (2014) Scyl1 scaffolds class II Arfs to specific subcomplexes of coatomer through the -COP appendage domain. J. Cell. Sci. 127: 1454–1463

Hammarén HM, Ungureanu D, Grisouard J, Skoda RC, Hubbard SR & Silvennoinen O (2015) ATP binding to the pseudokinase domain of JAK2 is critical for pathogenic activation. Proc Natl Acad Sci USA 112: 4642–4647

Hartman T, Stead K, Koshland D & Guacci V (2000) Pds5p Is an Essential Chromosomal Protein Required for Both Sister Chromatid Cohesion and Condensation in Saccharomyces cerevisiae. J. Cell Biol. 151: 613–626

Hopper AK (2013) Transfer RNA post-transcriptional processing, turnover, and subcellular dynamics in the yeast Saccharomyces cerevisiae. Genetics 194: 43–67

Horazdovsky BF, Busch GR & Emr SD (1994) VPS21 encodes a rab5-like GTP binding protein that is required for the sorting of yeast vacuolar proteins. The EMBO Journal 13: 1297–1309

Huh W-K, Falvo JV, Gerke LC, Carroll AS, Howson RW, Weissman JS & O’Shea EK (2003) Global analysis of protein localization in budding yeast. Nature 425: 686–691

Ioannou MG & Summerfeldt N (2015) Scyl1 scaffolds class II Arfs to selective subcomplexes of coatomer via the γ-COP appendage domain. … of COPI trafficking

Izaurralde E, Kutay U, Kobbe von C, Mattaj IW & Görlich D (1997) The asymmetric distribution of the constituents of the Ran system is essential for transport into and out of the nucleus. The EMBO Journal 16: 6535–6547

Johnson LM, Bankaitis VA & Emr SD (1987) Distinct sequence determinants direct intracellular sorting and modification of a yeast vacuolar protease. Cell 48: 875–885

Johnstone AD, Mullen RT & Mangroo D (2011) ArabidopsisAt2g40730 encodes a cytoplasmic protein involved in nuclear tRNA export. Botany 89: 175–190

Kato M, Yano K-I, Morotomi-Yano K, Saito H & Miki Y (2002) Identification and Characterization of the Human Protein Kinase-like Gene NTKL: Mitosis-Specific Centrosomal Localization of an Alternatively Spliced Isoform. Genomics 79: 760–767

Kitamoto K, Yoshizawa K, Ohsumi Y & Anraku Y (1988) Dynamic aspects of vacuolar and cytosolic amino acid pools of Saccharomyces cerevisiae. J. Bacteriol. 170: 2683–2686

Majcher V, Goode A, James V & Layfield R (2015) Autophagy receptor defects and ALS-FTLD. Mol. Cell. Neurosci. 66: 43–52

Marcusson EG, Horazdovsky BF, Cereghino JL, Gharakhanian E & Emr SD (1994) The sorting receptor for yeast vacuolar carboxypeptidase Y is encoded by the VPS10 gene. Cell 77: 579–586

Matsuyama A, Arai R, Yashiroda Y, Shirai A, Kamata A, Sekido S, Kobayashi Y, Hashimoto A, Hamamoto M, Hiraoka Y, Horinouchi S & Yoshida M (2006) ORFeome cloning and global analysis of protein localization in the fission yeast Schizosaccharomyces pombe. Nature Biotechnology 24: 841–847

McGuire AT & Mangroo D (2007) Cex1p is a novel cytoplasmic component of the Saccharomyces cerevisiae nuclear tRNA export machinery. The EMBO Journal 26: 288–300

McGuire AT & Mangroo D (2012) Cex1p facilitates Rna1p-mediated dissociation of the Los1p-tRNA-Gsp1p-GTP export complex. Traffic 13: 234–256

Murphy JM, Zhang Q, Young SN, Reese ML, Bailey FP, Eyers PA, Ungureanu D, Hammaren H, Silvennoinen O, Varghese LN, Chen K, Tripaydonis A, Jura N, Fukuda K, Qin J, Nimchuk Z, Mudgett MB, Elowe S, Gee CL, Liu L, et al (2014) A robust methodology to subclassify pseudokinases based on their nucleotide-binding properties. Biochem. J. 457: 323–334

Nozawa K, Ishitani R, Yoshihisa T, Sato M, Arisaka F, Kanamaru S, Dohmae N, Mangroo D, Senger B, Becker HD & Nureki O (2013) Crystal structure of Cex1p reveals the mechanism of tRNA trafficking between nucleus and cytoplasm. Nucl. Acids Res. 41: 3901–3914

Panizza S, Tanaka T, Hochwagen A, Eisenhaber F & Nasmyth K (2000) Pds5 cooperates with cohesin in maintaining sister chromatid cohesion. Current Biology 10: 1557–1564

Pelletier S, Gingras S, Howell S, Vogel P & Ihle JN (2012) An Early Onset Progressive Motor Neuron Disorder in Scyl1-Deficient Mice Is Associated with Mislocalization of TDP-43. Journal of Neuroscience 32: 16560–16573

Piper RC, Cooper AA, Yang H & Stevens TH (1995) VPS27 controls vacuolar and endocytic traffic through a prevacuolar compartment in Saccharomyces cerevisiae. J. Cell Biol. 131: 603–617

Prescianotto Baschong C & Riezman H (2002) Ordering of Compartments in the Yeast Endocytic Pathway. Traffic 3: 37–49

Richards SA, Carey KL & Macara IG (1997) Requirement of guanosine triphosphate-bound ran for signal-mediated nuclear protein export. Science 276: 1842–1844

Richardson BC, Halaby SL, Gustafson MA & Fromme JC (2016) The Sec7 N-terminal regulatory domains facilitate membrane-proximal activation of the Arf1 GTPase. Elife 5: 11711

Robinson JS, Klionsky DJ, Banta LM & Emr SD (1988) Protein sorting in Saccharomyces cerevisiae: isolation of mutants defective in the delivery and processing of multiple vacuolar hydrolases. Mol. Cell. Biol. 8: 4936–4948

Schmidt WM1, Kraus C, Höger H, Hochmeister S, Oberndorfer F, Branka M, Bingemann S, Lassmann H, Müller M, Macedo-Souza LI, Vainzof M, Zatz M, Reis A, Bittner RE. (2007) Mutation in the Scyl1 gene encoding amino-terminal kinase-like protein causes a recessive form of spinocerebellar neurodegeneration. EMBO Rep. Jul. 8(7):691–7.

Schmidt WM, Rutledge SL, Schüle R, Mayerhofer B, Züchner S, Boltshauser E & Bittner RE (2015) Disruptive SCYL1 Mutations Underlie a Syndrome Characterized by Recurrent Episodes of Liver Failure, Peripheral Neuropathy, Cerebellar Atrophy, and Ataxia. Am. J. Hum. Genet. 97: 855–861

Shaheen HH & Hopper AK (2005) Retrograde movement of tRNAs from the cytoplasm to the nucleus in Saccharomyces cerevisiae. Proceedings of the National Academy of Sciences 102: 11290–11295

Simos G, Segref A, Fasiolo F, Hellmuth K, Shevchenko A, Mann M & Hurt EC (1996) The yeast protein Arc1p binds to tRNA and functions as a cofactor for the methionyl-and glutamyl-tRNA synthetases. The EMBO Journal 15: 5437–5448

Stanford DR, Whitney ML, Hurto RL, Eisaman DM, Shen W-C & Hopper AK (2004) Division of labor among the yeast Sol proteins implicated in tRNA nuclear export and carbohydrate metabolism. Genetics 168: 117–127

Stead K, Aguilar C, Hartman T, Drexel M, Meluh P & Guacci V (2003) Pds5p regulates the maintenance of sister chromatid cohesion and is sumoylated to promote the dissolution of cohesion. J. Cell Biol. 163: 729–741

Steiner-Mosonyi M, Leslie DM, Dehghani H, Aitchison JD & Mangroo D (2003) Utp8p Is an Essential Intranuclear Component of the Nuclear tRNA Export Machinery of Saccharomyces cerevisiae. Journal of Biological Chemistry 278: 32236–32245

Stevens T, Esmon B & Schekman R (1982) Early stages in the yeast secretory pathway are required for transport of carboxypeptidase Y to the vacuole. Cell 30: 439–448

Shibuya A, Margulis N, Christiano R, Walther TC & Barlowe C (2015) The Erv41-Erv46 complex serves as a retrograde receptor to retrieve escaped ER proteins. J. Cell Biol. 208: 197–209

Sullivan BM, Harrison-Lavoie KJ, Marshansky V, Lin HY, Kehrl JH, Ausiello DA, Brown D & Druey KM (2000) RGS4 and RGS2 bind coatomer and inhibit COPI association with Golgi membranes and intracellular transport. Mol. Biol. Cell 11: 3155–3168

Usaj M, Tan Y, Wang W, VanderSluis B, Zou A, Myers CL, Costanzo M, Andrews B & Boone C (2017) TheCellMap.org: A Web-Accessible Database for Visualizing and Mining the Global Yeast Genetic Interaction Network. G3 (Bethesda) 7: 1539–1549

Vida TA & Emr SD (1995) A new vital stain for visualizing vacuolar membrane dynamics and endocytosis in yeast. J. Cell Biol. 128: 779–792

Walch-Solimena C & Novick P (1999) The yeast phosphatidylinositol-4-OH kinase Pik1 regulates secretion at the Golgi. Nature Cell Biology 1: 523–525

Wang K, Yang Z, Liu X, Mao K, Nair U & Klionsky DJ (2012) Phosphatidylinositol 4-kinases are required for autophagic membrane trafficking. J. Biol. Chem. 287: 37964–37972

Yoshihisa T, Yunoki-Esaki K, Ohshima C, Tanaka N & Endo T (2003) Possibility of cytoplasmic pre-tRNA splicing: the yeast tRNA splicing endonuclease mainly localizes on the mitochondria. Mol. Biol. Cell 14: 3266–3279

Yoshimura SH & Hirano T (2016) HEAT repeats - versatile arrays of amphiphilic helices working in crowded environments? J Cell Sci 129: 3963–3970

Zabezhinsky D, Slobodin B, Rapaport D, Gerst JE (2016) An Essential Role for COPI in mRNA Localization to Mitochondria and Mitochondrial Function. Cell Rep.15(3):540–549

